# APOBEC3 mutagenesis drives therapy resistance in breast cancer

**DOI:** 10.1101/2024.04.29.591453

**Authors:** Avantika Gupta, Andrea Gazzo, Pier Selenica, Anton Safonov, Fresia Pareja, Edaise M. da Silva, David N. Brown, Yingjie Zhu, Juber Patel, Juan Blanco-Heredia, Bojana Stefanovska, Michael A. Carpenter, Xin Pei, Denise Frosina, Achim A. Jungbluth, Marc Ladanyi, Giuseppe Curigliano, Britta Weigelt, Nadeem Riaz, Simon N. Powell, Pedram Razavi, Reuben S. Harris, Jorge S. Reis-Filho, Antonio Marra, Sarat Chandarlapaty

**Author notes:** Corresponding authors: Antonio Marra and Sarat Chandarlapaty. These authors contributed equally to this article (Avantika Gupta, Andrea Gazzo). Early Drug Development for Innovative Therapies, European Institute of Oncology IRCSS, Milan, Italy.

## Abstract

Acquired genetic alterations commonly drive resistance to endocrine and targeted therapies in metastatic breast cancer^1–7^, however the underlying processes engendering these diverse alterations are largely uncharacterized. To identify the mutational processes operant in breast cancer and their impact on clinical outcomes, we utilized a well-annotated cohort of 3,880 patient samples with paired tumor-normal sequencing data. The mutational signatures associated with apolipoprotein B mRNA-editing enzyme catalytic polypeptide-like 3 (APOBEC3) enzymes were highly prevalent and enriched in post-treatment compared to treatment-naïve hormone receptor-positive (HR+) cancers. APOBEC3 mutational signatures were independently associated with shorter progression-free survival on antiestrogen plus CDK4/6 inhibitor combination therapy in patients with HR+ metastatic breast cancer. Whole genome sequencing (WGS) of breast cancer models and selected paired primary-metastatic samples demonstrated that active APOBEC3 mutagenesis promoted resistance to both endocrine and targeted therapies through characteristic alterations such as *RB1* loss-of-function mutations. Evidence of APOBEC3 activity in pre-treatment samples illustrated a pervasive role for this mutational process in breast cancer evolution. The study reveals APOBEC3 mutagenesis to be a frequent mediator of therapy resistance in breast cancer and highlights its potential as a biomarker and target for overcoming resistance.

## MAIN

Both endocrine and targeted therapies are broadly utilized to treat breast cancer, however their potential for durable, long-term disease control is limited by tumor evolution and the development of acquired genetic alterations that promote drug resistance^8^. In estrogen receptor (ER)-positive cancers, acquired alterations in several genes including estrogen receptor (*ESR1*), neurofibromin (*NF1*)^9^, and v-erb-b2 avian erythroblastic leukemia viral oncogene homolog 2 (*ERBB2*, also known as HER2) are frequently observed as tumors develop resistance to antiestrogen therapy. Such acquired genetic alterations have also been found after development of resistance to targeted therapies including cyclin dependent kinase 4/6 (CDK4/6) inhibitors, HER2 inhibitors, and phosphoinositide 3-kinase (PI3K) inhibitors^4,10–12^. The widespread prevalence of these heterogeneous and acquired genetic alterations has proven challenging to overcome despite the development of several second-generation inhibitors such as selective estrogen receptor degraders (SERDs) and have highlighted the need to understand the underlying processes behind the genomic instability and capacity for tumor evolution among breast cancers.

Detailed analyses of cancer genomes have cataloged the presence of discrete mutational signatures that can be tumor-type specific, readily promote tumor evolution, and potentially predict rational treatment strategies^13–15^. In breast cancer, single base substitution (SBS) signatures associated with the activity of APOBEC3 enzymes (COSMIC SBS2 and SBS13) are highly prevalent suggesting the potential relevance for these mutational processes in the underlying genomic instability. Prior studies have linked high expression levels of specific APOBEC3 family enzymes to inferior outcomes and tamoxifen resistance in ER-positive tumors and xenograft models^16,17^. Moreover, APOBEC3 mutational signatures are reported to be enriched in metastatic compared to primary breast cancers^18,19^. Indeed, APOBEC3 mutagenesis has been recently proposed to be a contributor towards clonal evolution of metastatic ER-positive breast cancers treated with endocrine-based therapy^20^. These and other studies reveal a potential association of APOBEC3 mutagenesis with tumor progression and raise the possibility that these processes may be ongoing and contributory to treatment resistance. Herein, we report that APOBEC3 mutagenesis driven by APOBEC3A (A3A) and APOBEC3B (A3B) facilitates breast cancer evolution independent of treatment exposure, leading to resistance against a diverse range of therapeutic agents through the induction of APOBEC3-class alterations in characteristic resistance-associated genes. The results highlight the potential for APOBEC3 mutagenesis as a readily detectable biomarker for distinct resistance trajectories pointing to alternative approaches to target these evolvable cancers.

### APOBEC3 mutagenesis is highly prevalent in metastatic and treatment-resistant breast cancers

To characterize the mutational landscape of treatment-sensitive and -resistant breast cancers broadly, we leveraged a cohort including over 5,000 breast cancer samples that were previously subjected to paired tumor-normal sequencing by the MSK-IMPACT assay^21^. After excluding samples with sequencing-estimated low tumor purity (see **Methods**), 3,880 high-quality samples from 3,117 patients were analyzed (**Fig. 1a**). This cohort constitutes a large collection of genomically-profiled breast cancers coupled with detailed clinical annotation, including treatment and follow-up information, and is representative of the clinical diversity of the disease encompassing all subtypes, histologies, tumor grade and other clinical characteristics (**Supplementary Table 1**). To investigate the contribution of mutational processes to clinical characteristics and outcomes of breast cancers, we utilized the Single Multivariate Analysis (*SigMA*) tool^22^ to deconvolute mutational signatures in our study population. *SigMA* has been previously validated for assessment of homologous recombination deficiency (HRD) and other mutational processes in solid tumors^23–25^. To benchmark its validity and fidelity to evaluate APOBEC3 mutational signatures in breast cancers assessed using a targeted sequencing panel, we down-sampled publicly-available whole exome sequencing (WES) data, including primary breast cancers from The Cancer Genome Atlas (TCGA) (*n* = 1019) and metastatic breast cancers from *Bertucci et al.* (*n* = 617)^18^ as well as whole genome sequencing (WGS) data of primary breast cancers from *Nik-Zainal et al.* (*n* = 560)^26^, to the genomic footprint of the MSK-IMPACT panel. We computed the dominant mutational signatures (see **Methods**) in the down-sampled data with *SigMA* and compared with the dominant signatures called on the WES and WGS data using multiple analytic tools^27–29^ (**Fig. S1a**). *SigMA* displayed high sensitivity (0.9 – 0.93), specificity (0.79 – 0.84) and accuracy (0.82 – 0.86) in detecting APOBEC3 as the dominant signature (**Fig. S1b**). In addition to detecting APOBEC3 as the dominant signature, correlations for exposures calculated from simulated panels using *SigMA* and WES ranged from 0.75 – 0.79 (**Fig. S1c**). Finally, considering that *SigMA* requires ≥5 single nucleotide variants (SNVs) as input to assess mutational signatures from targeted sequencing, we found that only 0-4% of samples in the three analyzed WES/WGS datasets had <5 SNVs (**Fig. S1d**) indicating *SigMA’s* ability to process most samples from such datasets.

**Fig 1.**
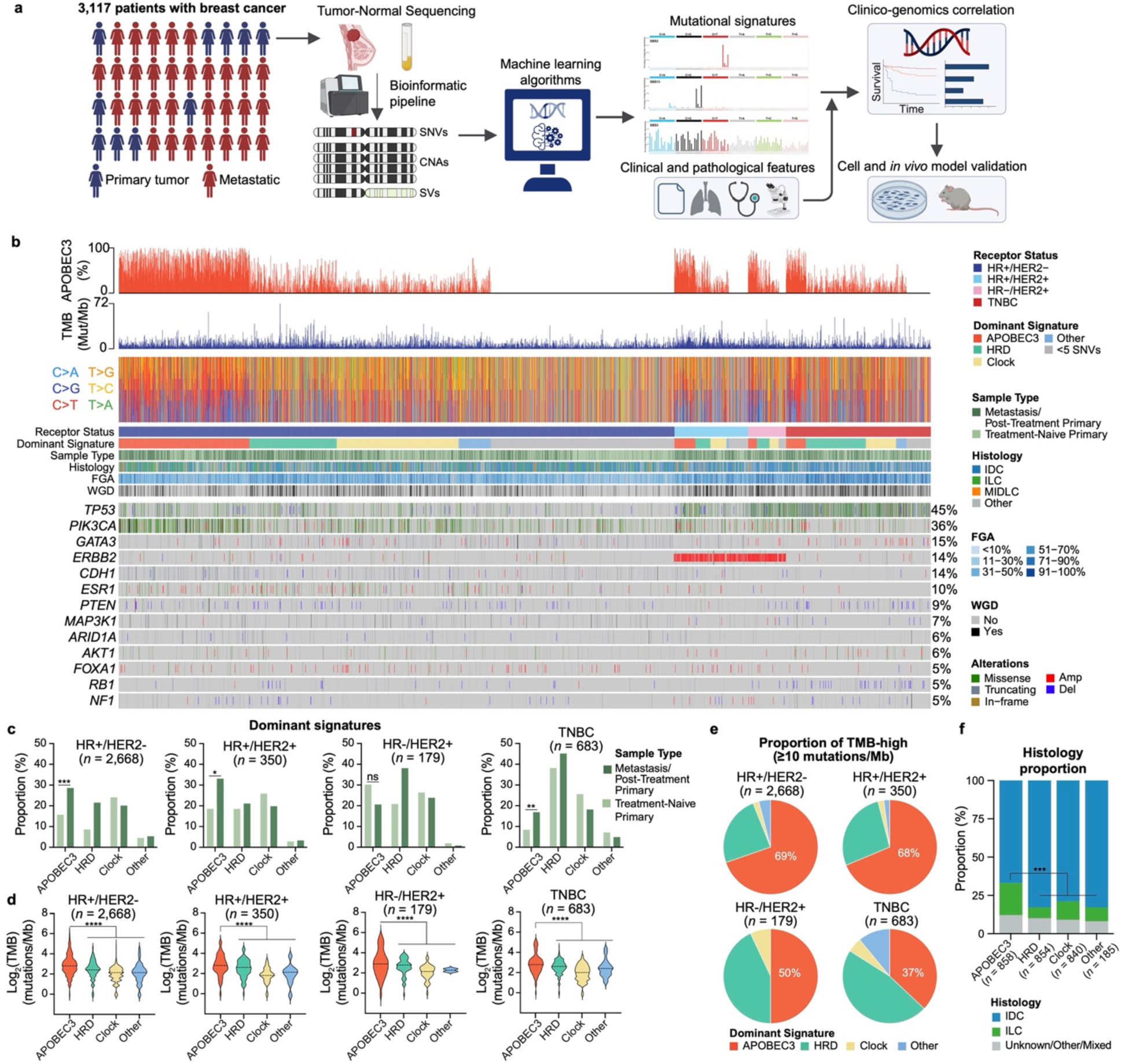
APOBEC3 mutational signatures are prevalent in breast cancers. **(a)** Schematic of the analysis pipeline of the MSK-IMPACT breast cancer cohort. **(b)** Summary of the genomic characteristics of the clinical cohort demonstrating percentage contribution of APOBEC3 mutational signature (first panel), TMB (second panel), SNV change (third panel), and OncoPrint of select genes in samples. **(c)** Bar plots displaying the proportion of samples with indicated dominant mutational signature categorized by sample type and receptor status. Groups were compared using the two-tailed Wilcoxon test: ns not significant, * *p*<0.05, ** *p*<0.01, *** *p*<0.001 **(d)** Violin plots representing the TMB in samples categorized by receptor status. Groups were compared to APOBEC3-dominant samples using the two-tailed Wilcoxon test: **** *p*<0.0001. **(e)** Proportion of TMB-high samples with different dominant mutational signatures categorized by receptor status. **(f)** Proportion of samples categorized by dominant mutational signature and histology. Groups were compared to APOBEC3-dominant samples using the two-tailed Wilcoxon test: *** *p*<0.001.

Examining the 3,880 tumor samples with *SigMA*, the most prevalent, dominant mutational process was APOBEC3, consistent with previous observations from WGS and WES studies^13,18^ (**Fig. 1b**). APOBEC3-dominant signature was found in 15.7% and 28.7% of primary and metastatic HR+/HER2-, 18.5% and 33.1% of primary and metastatic HR+/HER2+, 30.2% and 20.6% of primary and metastatic HR-/HER2+, and 8.4% and 16.9% of primary and metastatic triple-negative breast cancers (TNBC) (**Fig. 1c**), respectively. APOBEC3 exposures were significantly higher in HR+ and TNBC metastatic breast cancers than in unmatched primary tumors (*p*<0.0001 for HR+/HER2-, *p*<0.01 for HR+/HER2+ and TNBC, **Fig. S2a**), suggesting a possible relationship to poor clinical outcomes. We next assessed the relationship between APOBEC3 dominant signature and indicators of genomic instability, such as tumor mutational burden (TMB), fraction of genome altered (FGA) and whole genome doubling (WGD). APOBEC3-dominant HR+/HER2 tumors displayed lower median FGA compared to non-APOBEC3 tumors independent of sample type (**Fig. S2b**). A similar pattern was also found in metastatic TNBC. Conversely, APOBEC3-dominant HR+ tumors had higher FGA compared to primary breast cancers of the same subtype (HR+/HER2-0.5 vs 0.4, *p*<0.01, HR+/HER2+ 0.7 vs 0.5, *p*<0.05) suggesting a higher APOBEC3-mediated genomic instability in the metastatic setting. The proportion of WGD was lower in APOBEC3-dominant HR+/HER2-metastatic tumors (35% vs 45%, *p*=0.3, **Fig. S2c**). APOBEC3-dominant breast cancers exhibited significantly higher median TMB than those with other mutational processes (*p*<0.0001 for all comparisons, **Fig. 1d**), underscoring the distinct impact of APOBEC3 mutagenesis. For metastatic breast cancers with TMB≥10 mutations per megabase, representing the current indication for the tumor agnostic use of anti-PD-1 immunotherapy^30^, APOBEC3 constituted the dominant mutational process in the vast majority of HR+/HER2- and HR+/HER2+ tumors (69% and 68%, respectively, **Fig. 1e**). In terms of standard clinical characteristics, groups of APOBEC3 dominant and other dominant signatures were highly similar (**Supplementary Table 2**). However, in terms of tumor histology, invasive lobular breast cancers (ILC, *n* = 489) were more frequently associated with APOBEC3 as dominant mutational signature as compared to invasive ductal and other histology types of breast cancer regardless of the sample type (38% ILCs were APOBEC3-dominant vs 13% HRD, 22% Clock and 3.3% Other, *p*<0.001) (**Fig. 1f**), consistent with previous evidence^31^.

### APOBEC3 enzymes cause APOBEC3-context point mutations and other genomic alterations

Out of the eleven APOBEC family members, A3A and A3B have emerged as the major drivers of APOBEC3 mutagenesis^32–36^. To assess the expression of these proteins among breast cancers harboring APOBEC3 mutational signatures, we utilized immunohistochemistry (IHC) to stain 130 tissue samples using an A3A-specific antibody^37^ or an antibody that detects A3A/B/G^38^ (**Fig. 2a**). We detected weak to strong levels of protein expression in 17/20 APOBEC3-dominant samples using an A3A/B/G antibody and 8/20 samples using an A3A-specific antibody, consistent with the reportedly weaker expression of A3A^39,40^. We also observed a weak to moderate correlation between APOBEC3 signature exposure and IHC-based expression (Pearson’s correlation coefficient = 0.41, *p*<0.01 for HR+/HER2-samples, **Fig. S3a**). Given the known cell cycle-dependent or episodic expression of APOBEC3 proteins^41–43^, and because mutational signatures reflect accumulation of genomic changes over the entire lifetime of a cell, these findings cannot establish whether high level of protein expression is necessary at the time of sample collection for signature manifestation. Our findings suggest that mutational signatures might be a more robust method to detect tumors with APOBEC3 activity.

**Fig 2.**
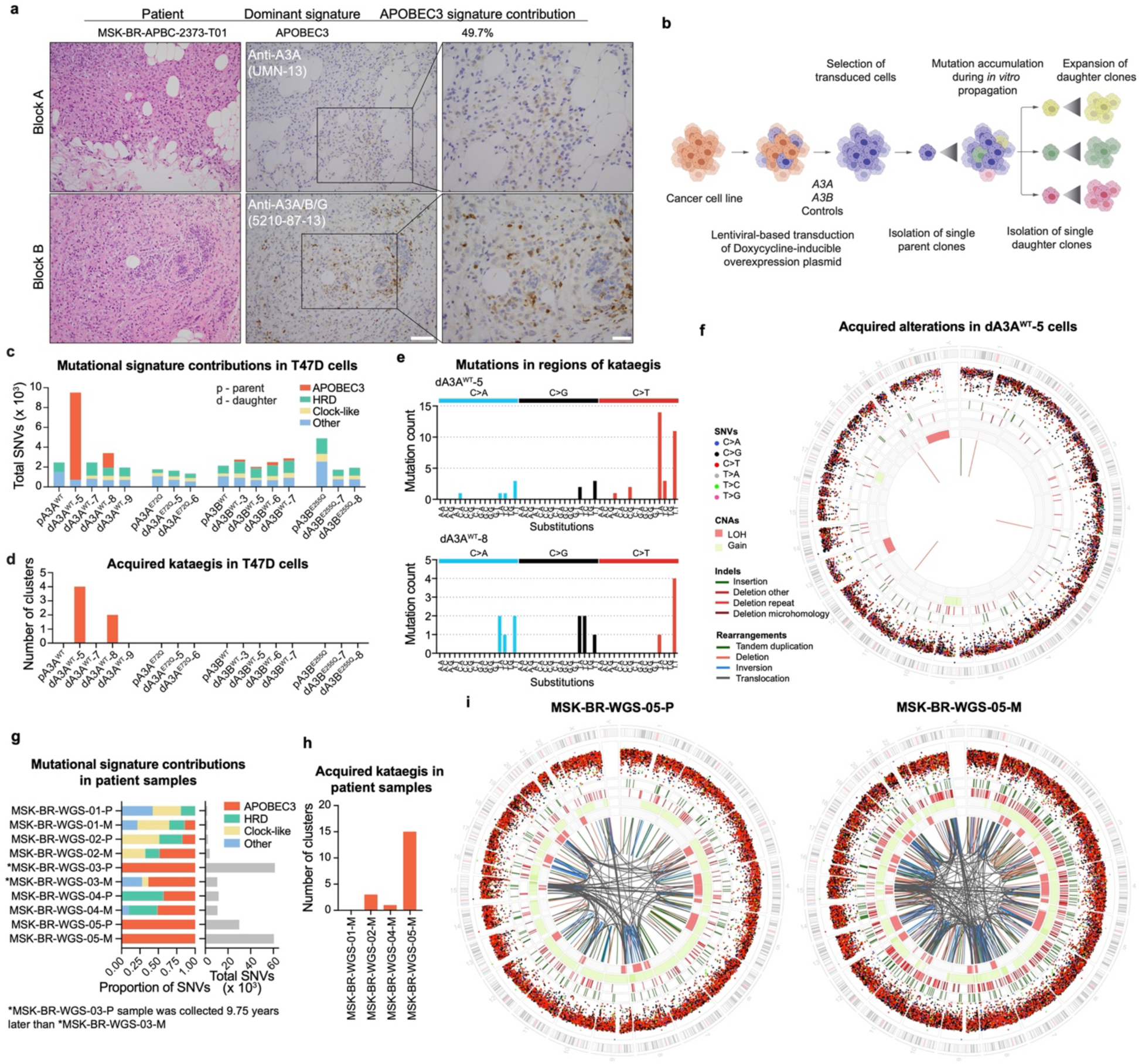
APOBE3 enzymes induce APOBEC3 mutational processes. **(a)** H&E and immunohistochemical images displaying A3A and A3A/B/G staining in an APOBEC3-dominant patient sample. Scale bar = 50 µm (middle panel), and 20 µm (right panel). **(b)** Schematic of the experimental design to investigate ongoing mutational processes in cells overexpressing WT A3A, A3B or their catalytic mutant controls. **(c)** Mutational signature contribution of acquired SNVs in the indicated samples. **(d)** Bar plots representing number of clusters with acquired regions of kataegis in samples from (c). **(e)** Substitution profile of dA3A^WT^-5 and dA3A^WT^-8 in the kataegis regions. **(f)** Circos plot representing acquired SNVs, indels, CNAs and structural rearrangements in dA3A^WT^-5 cells. **(g)** Mutational signature contribution and number of SNVs from WGS of five paired primary/metastatic patient samples. **(h)** Bar plots representing number of clusters with acquired regions of kataegis in the indicated metastatic patient samples. **(i)** Circos plots of samples MSK-BR-WGS-05-P and MSK-BR-WGS-05-M.

To directly evaluate if A3A or A3B can cause APOBEC3 mutagenesis in breast cancer cells, we overexpressed HA-tagged wild-type (WT) enzymes and their catalytically inactive mutants (A3A^E72Q^ or A3B^E255Q^) in ER-positive breast cancer cells lacking endogenous APOBEC3 activity^44^ using a doxycycline-inducible system (**Fig. S3b**). We observed *in vitro* DNA deaminase activity with overexpression of both WT enzymes but not with the mutant controls. To assess the acquired alterations upon A3A or A3B overexpression, we isolated single cell parent clones and exposed them to doxycycline for 72 days to drive mutagenesis followed by expansion of multiple single cell daughter clones (**Fig. 2b**). Interestingly, two out of four A3A WT expressing daughter clones lost protein expression and deaminase activity (**Fig. S3c**). Similar negative selection against A3A expression has been reported earlier^32,37^. We performed WGS of parent and daughter clones and analyzed acquired alterations. The lack of endogenous APOBEC3 activity in the model was confirmed by the absence of acquired APOBEC3 signature in the parental cells before doxycycline treatment (**Fig. 2c**). Among the daughter cells that maintained protein expression, A3A and A3B WT cells accumulated APOBEC3-context mutations (41.3 – 92.5% of the total SNVs in A3A and 6.7 – 12.5% in A3B), whereas the mutant cells did not (**Fig. 2c**). The daughter cells that lost A3A expression and activity did not display any acquired APOBEC3 exposure further confirming the necessity of cytidine deaminase activity for signature accumulation. As an additional confirmation of the proficiency of *SigMA* to detect APOBEC3 signatures in a controlled experiment, we simulated the WGS of cell lines to the MSK-IMPACT panel and used *SigMA* to detect mutational signatures. Among the samples with ≥5 SNVs, *SigMA* successfully classified dA3A^WT^-5 as APOBEC3-dominant, confirming the high performance obtained using the WES and WGS datasets (**Fig. S3d**). The WT cells exhibited a higher TMB compared to the catalytic mutant controls (mean TMB 7.86 for A3A^WT^ vs 2.75 for A3A^E72Q^, 4.49 for A3B^WT^ vs 3.36 for A3B^E255Q^, **Fig. S3e**). Cells with APOBEC3 as the dominant signature, APOBEC3-positive cells, also accumulated increased clustered mutations characterized as kataegis^45^ and omikli^46^, that have been previously linked to APOBEC3 activity (**Figs. 2d** and **S3f**). The majority of mutations in the regions of kataegis were substitutions in the APOBEC3 enzyme recognized T<u>C</u>N motifs (**Fig. 2e**). In addition to single nucleotide changes, comparative analyses revealed other acquired genomic alterations including insertions and deletions (indels), copy number alterations (CNAs) and structural variations (SVs) (**Fig. 2f**). To assess for such non-SNV alterations in the clinical cohort, we performed WGS of five pairs of primary and metastatic patient samples (**Fig. 2g**). These samples reflect breast cancers with low to high APOBEC3 signatures, some with acquired APOBEC3 signature in metastatic samples indicating APOBEC3 mutagenic activity at the time of treatment resistance. As an example, we present the progressive evolution of a HR+/HER2-tumor to 100% APOBEC3 exposure over more than 20 years of clinical history and therapeutic pressure of several lines of treatment, including endocrine therapies, CDK4/6 inhibitors, and PI3K-pathway inhibitors (paired samples from MSK-BR-WGS-03, **Fig. S3g**). Similar to our findings in cell line models, APOBEC3-positive samples acquired clustered mutations characterized as kataegis (**Fig. 2h**). APOBEC3-positive samples also exhibited substantial genomic alterations including indels, CNAs, SVs, and demonstrate evidence of chromothripsis, another hallmark process that has been associated with APOBEC3 activity^23,47^ (**Fig. 2i**). This phenomenon characterized by numerous clustered chromosomal rearrangements in a single event causing complex genomic alterations in specific chromosomal regions, is notably observed in sample MSK-BR-WGS-05-M on chromosomes 15, 8, and X. Given the similarities with APOBEC3-positive cells, we conclude that overexpression of A3A and A3B enzymes phenocopy characteristics of APOBEC3-driven tumors in a deamination-dependent manner.

### APOBEC3 activity promotes therapy resistance

To investigate the relationship between APOBEC3 signature and therapy resistance, we leveraged the detailed clinical annotations linked to our clinical cohort (**Fig. 1a**) and assessed the outcomes of APOBEC3-dominant cases on different forms of therapy. To avoid inconsistencies related to changes of APOBEC3 mutagenesis over time, only patients for whom the sequenced biopsy was acquired directly prior to the first line of the indicated treatment (+/-30 days) were included in the analysis. In HR+/HER2-metastatic breast cancers treated with endocrine therapy (ET) as single agent on first line of therapy (*n* = 111), APOBEC3-dominant tumors were associated with numerically shorter median progression-free survival (PFS) compared to tumors with other dominant signatures (8.6 vs 15.6 months, HR 1.4, 95% CI 0.9 – 2.2, *p*=0.1, **Fig. 3a**). When comparing outcomes on first-line therapy with CDK4/6 inhibitor plus ET (*n* = 549), APOBEC3- and HRD-dominant metastatic breast cancers were independently associated with lower median PFS compared to other cancers regardless of ET partner and line of therapy in the metastatic setting (HR 1.5, 95% CI 1.2 - 1.8, *p*<0.001 and HR 1.8, 95% CI 1.4 - 2.2, *p*<0.001, respectively, **Fig. 3b**), suggesting that an increased genomic instability confers resistance to standard frontline therapy in HR+/HER2-advanced breast cancer.

**Fig 3.**
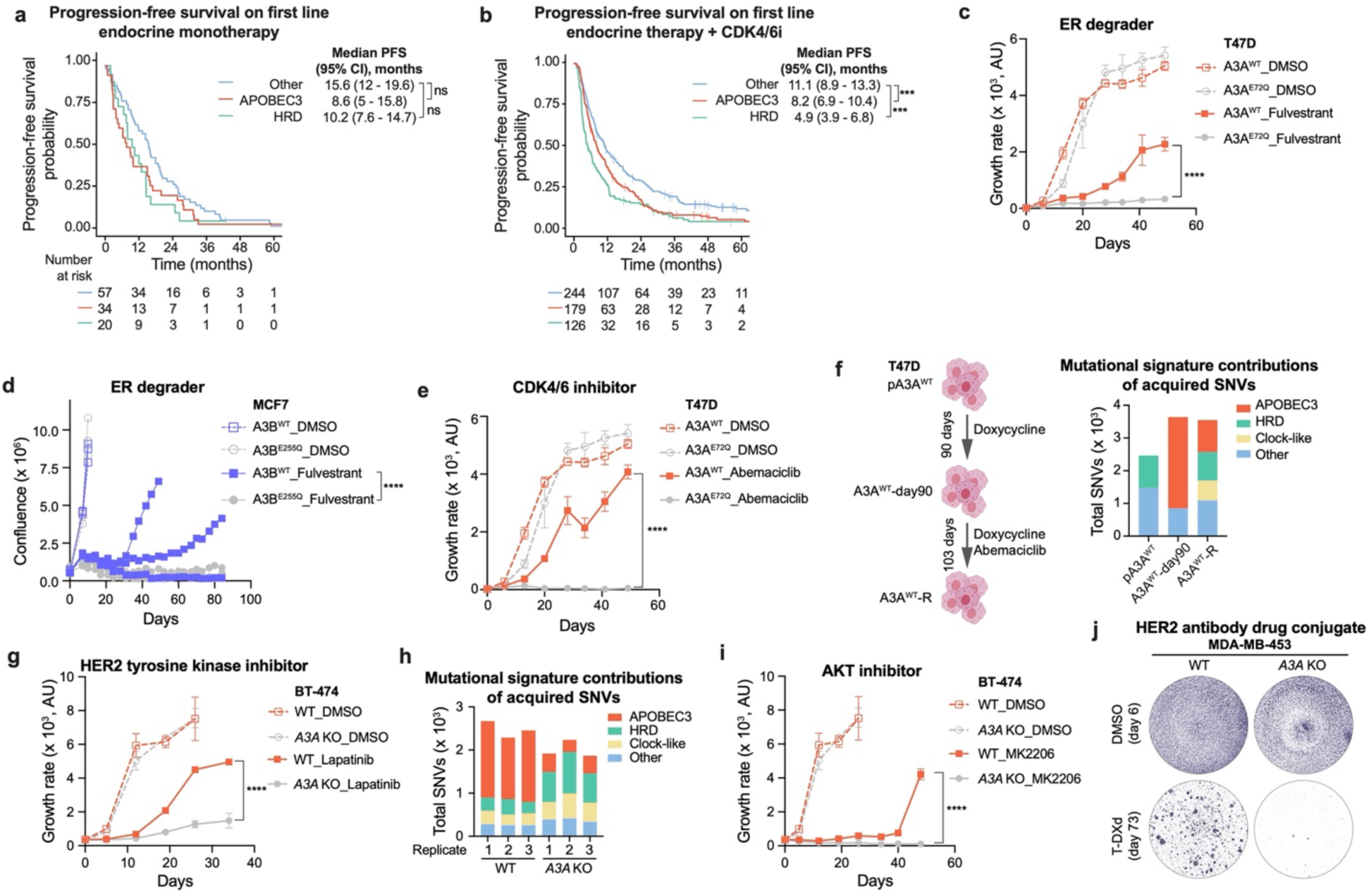
APOBEC3 mutagenesis promotes therapeutic resistance in breast cancers. **(a, b)** Kaplan-Meier curves displaying progression-free survival probability of patients with HR+/HER2-metastatic breast cancers treated with endocrine therapy **(a)** as single agent or **(b)** in combination with CDK4/6i. The patients are categorized according to the dominant mutational signatures of the biopsy obtained prior to treatment start. The groups were compared using log-rank test: ns not significant, *** *p*<0.001. **(c)** Growth curves of T47D A3A^WT^ and A3A^E72Q^ cells treated with DMSO or fulvestrant (10 nM). Data are represented as mean ± SD of three replicates. The groups were compared using two-way ANOVA test: **** *p*<0.0001. AU is abstract unit. **(d)** Growth curves of MCF7 A3B^WT^ and A3B^E255Q^ cells treated with DMSO or fulvestrant (10 nM). Data are represented as individual replicates (*n* = 3). The groups were compared using two-tailed Mann-Whitney U test: **** *p*<0.0001. **(e)** Growth curves of T47D A3A^WT^ and A3A^E72Q^ cells treated with DMSO or abemaciclib (500 nM). Data are represented as mean ± SD of three replicates. The groups were compared using two-way ANOVA test: **** *p*<0.0001. AU is abstract unit. **(f)** Schematic showing the timeline of generation of abemaciclib-resistant T47D A3A^WT^ (A3A^WT^-R) cells (left panel). Mutational signature contribution of acquired SNVs in the indicated samples (right panel). **(g)** Growth curves of BT-474 WT and *A3A* KO cells treated with DMSO or lapatinib (20 nM). Data are represented as mean ± SD of three replicates. The groups were compared using two-way ANOVA test: **** *p*<0.0001. AU is abstract unit. **(h)** Mutational signature contribution of acquired SNVs in the indicated samples. **(i)** Growth curves of BT-474 WT and *A3A* KO cells treated with DMSO or MK2206 (100 nM). Data are represented as mean ± SD of three replicates. The groups were compared using two-way ANOVA test: **** *p*<0.0001. AU is abstract unit. **(j)** Crystal violet staining of MDA-MB-453 WT and *A3A* KO cells treated with DMSO (for 6 days) or T-DXd (100 ng/mL, for 73 days). Images are representative of *n* = 3 replicates.

To assess the causal relationship between therapy resistance and APOBEC3 mutagenesis, we employed our long-term doxycycline-treated APOBEC3-positive and APOBEC3-negative models. The growth rate of APOBEC3-positive A3A^WT^ and APOBEC3-negative A3A^E72Q^ cells was comparable under DMSO treatment (**Fig. 3c**). When exposed to the SERD fulvestrant, A3A^WT^ cells acquired resistance significantly faster than A3A^E72Q^ cells (*p*<0.0001). We also assessed the effect of the weaker mutator A3B in influencing resistance in a similar experiment by analyzing the area occupied by cells. All A3B^WT^ and A3B^E255Q^ replicates grew similarly under DMSO. When grown under selection, 2/3 replicates of A3B^WT^ cells acquired resistance to fulvestrant, whereas none of the three A3B^E255Q^ replicates developed resistance after 84 days of continuous drug exposure (*p*<0.0001, **Fig. 3d**). A3A WT overexpression also led to a selective growth advantage of A3A^WT^ cells on treatment with the CDK4/6 inhibitors abemaciclib (*p*<0.0001, **Fig. 3e**) and palbociclib (*p*<0.0001, **Fig. S4a**). Importantly, based on analysis of the acquired alterations using WGS, A3A^WT^ cells continued to accumulate APOBEC3 signature mutations during resistance acquisition (**Fig. 3f**) further implying a role of APOBEC3 activity in driving resistance.

We next tested the effect of endogenous APOBEC3 enzymes in facilitating resistance. For this, we utilized the HER2+ models with endogenously active APOBEC3 mutagenesis that is predominantly driven by A3A^34^. Similar to our ER-positive models, the growth rates of APOBEC3-positive BT-474 WT cells and APOBEC3-negative *A3A* knockout (KO) cells were comparable under DMSO treatment (**Fig. 3g**). The WT cells acquired resistance to the tyrosine kinase inhibitor (TKI) lapatinib significantly faster than the *A3A* KO cells (*p*<0.0001). As with the continued APOBEC3 mutagenic activity observed in our ectopic overexpression system, BT-474 cells maintained endogenous APOBEC3 activity during treatment pressure (**Fig. 3h**) as confirmed by WGS of pre- and post-treatment cells. We also observed a significant growth advantage of MDA-MB-453 WT cells to TKIs lapatinib and neratinib compared to *A3A* KO cells (*p*<0.0001, **Fig. S4b-d**). We examined other anti-HER2 therapies including an inhibitor of the downstream target AKT kinase (AKT), MK2206, or the antibody drug conjugate, trastuzumab deruxtecan (T-DXd). BT-474 WT cells selectively gained resistance to MK2206 (*p*<0.0001, **Fig. 3i**) and MDA-MB-453 cells to T-DXd (**Fig. 3j**) compared to KO cells. Overall, our data demonstrate that APOBEC3 activity can readily promote therapy resistance in different subtypes of breast cancers and to a diverse range of therapeutic agents.

### Mechanisms of APOBEC3-mediated resistance

To mechanistically understand how APOBEC3 mutagenesis drives resistance in breast cancers, we sought to assess whether any of the alterations in genes linked to therapy resistance might be specifically brought about by APOBEC3 mutagenesis. We first conducted a gene-enrichment analysis using all samples within the MSK-IMPACT breast cancer cohort (*n* = 3,880, **Fig. 1b**). To address potential biases stemming from the APOBEC3-induced hypermutator phenotype, a permutation test was applied (see **Methods**). A statistically significant enrichment of oncogenic mutations affecting *PIK3CA*, *CDH1*, and *KMT2C* (*q*<0.1, mutated in >5% samples) was found in metastatic/post-treatment HR+/HER2-tumors bearing dominant APOBEC3 signatures (**Fig. 4a**). Similarly, APOBEC3-dominant, treatment-naïve HR+/HER2-or all TNBC samples also exhibited an enrichment of *PIK3CA* variants in contrast to non-APOBEC3 dominant tumors (**Fig. S5a-c**). Oncogenic mutations in other resistance-associated genes such as *NF1* and *ZFHX3* were also enriched in APOBEC3-dominant samples, albeit at a lower frequency (2.8% for *NF1*, and 1.1% for *ZFHX3*). We next examined acquired alterations in patients with HR+/HER2-breast cancer for which multiple tumor samples had been collected over time (patients with 2 samples, *n* = 449, patients with ≥3 samples, *n* = 43), focusing on APOBEC3-context mutations in samples with evidence of APOBEC3 activity. After *de novo* genotyping of somatic mutations found in each sample from the same patient (see **Methods**), we observed an enrichment of APOBEC3-context, acquired alterations in genes encoding transcription factors previously linked to endocrine therapy resistance, such as *ARID1A* and *ZFHX3*^48,49^. These alterations were enriched in treatment-resistant tumors with dominant APOBEC3 signature compared to non-APOBEC3 dominant samples (12.7% vs 3%, *q*<0.1 for *ARID1A*, 9% vs 1.4%, *q*<0.1 for *ZFHX3*) (**Fig. 4b**). The proportions of acquired alterations in key genes of the PI3K/Akt pathway, including *PIK3CA* (13% vs 6%, *q*>0.1) and *PTEN* (11% vs 6%, *q*>0.1), and other resistance-linked genes like *KMT2C* (9% vs 2%, *q*>0.1) were also numerically higher in APOBEC3-dominant, therapy-resistant samples compared to non-APOBEC3 dominant samples. We found that the proportion of APOBEC3-context mutations exclusive (acquired) to samples collected at later stages in the patient’s history was notably higher than those shared with early or treatment-naïve samples from the same patient (51.9% vs 32.3%, *p*<0.001, **Fig. 4c**) suggesting an active role of APOBEC3 mutagenesis in driving resistance-linked mutations described above.

**Fig 4.**
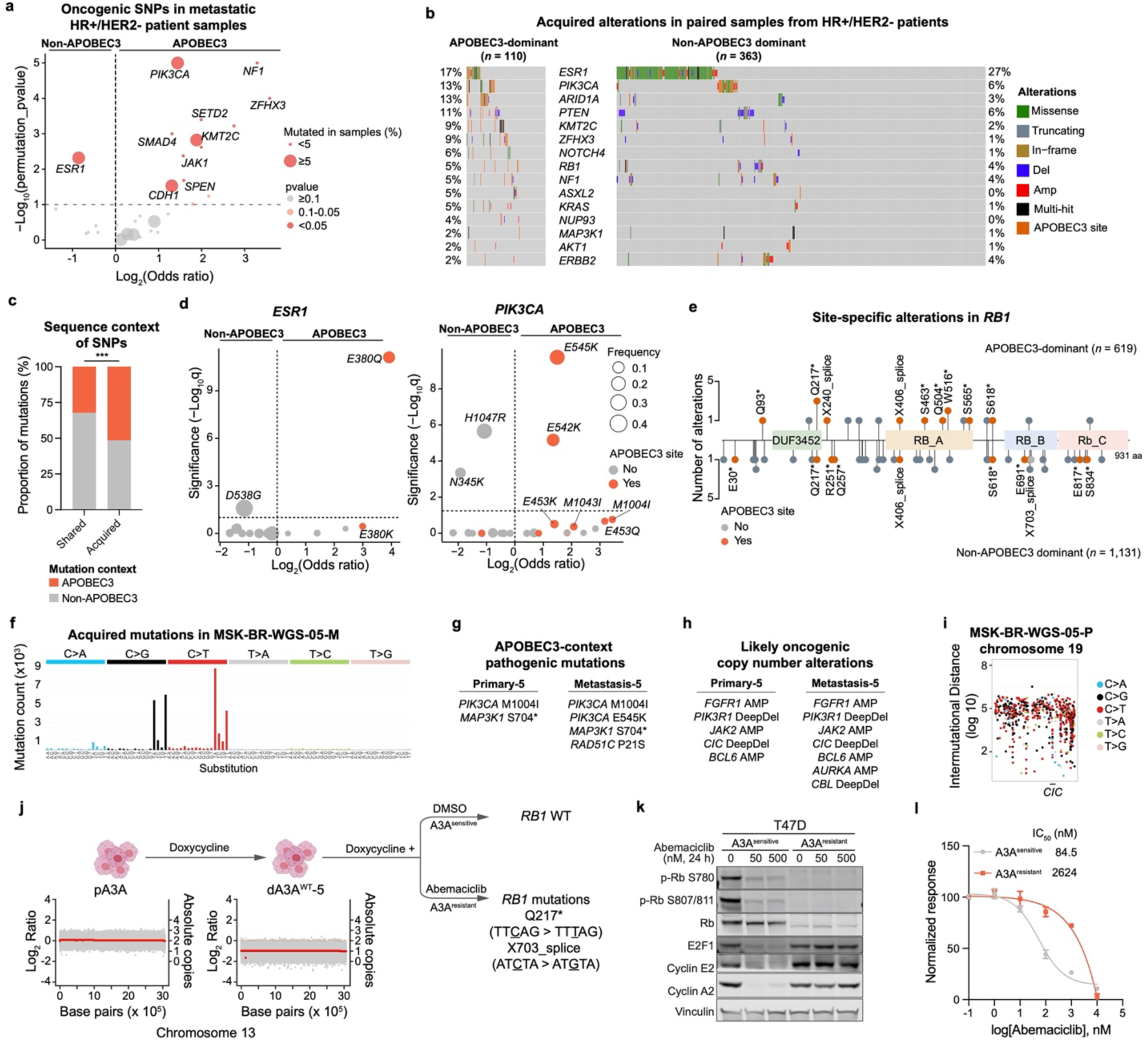
APOBEC3-class alterations drive therapeutic resistance in breast cancers. **(a)** Volcano plot displaying enrichment of genes in metastatic HR+/HER2-samples categorized according to the dominant mutational signature. **(b)** OncoPrint of acquired alterations in HR+/HER2-tumor samples categorized according to the dominant mutational signature. **(c)** Barplots representing proportion of shared or acquired SNPs categorized as APOBEC3-context and non-APOBEC3 context SNPs. The groups were compared using the Fisher’s exact test: *** *p*<0.001. **(d)** Volcano plots depicting site-specific enrichment of *ESR1* (left panel) and *PIK3CA* (right panel) categorized according to the dominant mutational signature. **(e)** Site-specific enrichment of alterations in *RB1* gene in HR+/HER2-tumor samples. **(f)** Mutation spectrum of acquired SNVs in MSK-BR-WGS-05-M. **(g)** Pathogenic mutations with APOBEC3-context substitutions in samples MSK-BR-WGS-05-P and MSK-BR-WGS-05-M. **(h)** Likely oncogenic CNAs in samples MSK-BR-WGS-05-P and MSK-BR-WGS-05-M. **(i)** Rainfall plot displaying intermutational distance between SNVs in chromosome 19 of sample MSK-BR-WGS-05-P. **(j)** FACETS plot showing LOH of chromosome 13 in dA3A^WT^-5 and acquired SNVs in *RB1* after DMSO treatment (A3A^sensitive^) or abemaciclib (1 µM) treatment (A3A^resistant^). **(k)** Immunoblots displaying changes in cell cycle regulatory proteins in A3A^sensitive^ and A3A^resistant^ cells treated with DMSO or indicated doses of abemaciclib for 24 h. Vinculin was used as a loading control. Immunoblots are representative of *n* = 3 independent experiments. **(l)** Inhibition of proliferation of A3A^sensitive^ and A3A^resistant^ cells treated with increasing concentrations of abemaciclib. Data are represented as mean ± SD of three replicates and are normalized to the DMSO control.

Alterations in *ESR1* comprise the most frequent class of acquired resistance alterations in breast cancer. While studying the association of APOBEC3 mutagenesis with genetic alterations in our paired analysis of HR+/HER2-samples, we observed that 17% of APOBEC3-dominant samples presented with *ESR1* alterations compared to 27% non-APOBEC3 dominant samples (*q*<0.1, **Fig. 4b**). To explore this further, we examined site-specific gene enrichment of all genes included in the MSK-IMPACT panel that were previously linked to endocrine therapy resistance (**Fig. S6**). Strikingly, *ESR1* E380Q mutations, an APOBEC3-context substitution, were specifically enriched in APOBEC3-dominant, HR+/HER2-post-treatment samples (*q*<0.1, **Fig. 4d**). By contrast, highly activating *ESR1* mutations in the loop between helix 11 and helix 12, (mutations in L536X, Y537X, and D538X, none of which are APOBEC3 context substitutions) were rare in APOBEC3 dominant samples (8%) suggesting that APOBEC3-dominant tumors might not dominantly reactivate ER as the mechanism of endocrine resistance. Among other common mutations, *PIK3CA* hotspot mutations, such as E545K, E542K, and E453Q, which are APOBEC3-context mutations, were predominantly detected in APOBEC3-dominant HR+/HER2-post-treatment samples (*q*<0.1, **Fig. 4d**), confirming the enrichment of *PIK3CA* helical domain mutations in APOBEC3-dominant breast cancers^50^. These findings suggest that APOBEC3 contributes to site-specific resistant-associated alterations, but not necessarily all resistant-causing changes in a gene, also exemplified by those observed for the tumor suppressor *RB1* (**Fig. 4e**).

As a salient example of involvement of APOBEC3 mutagenesis in acquired resistance to endocrine therapy, we explored in depth the WGS of patient MSK-BR-WGS-05 (**Fig. 2g**) who received endocrine therapy including the selective ER modulator tamoxifen and the aromatase inhibitor letrozole. In this case, 95% of acquired mutations were assigned to APOBEC3 signature (**Fig. 4f**) implying that APOBEC3 mutational processes were active at some point during progression and resistance development. The cancer specifically acquired APOBEC3-context pathogenic mutations in *PIK3CA* (E545K) and *RAD51C* (P21S) (**Fig. 4g**). In terms of CNAs, the cancer maintained the *FGFR1* amplification and *PIK3R1* deep deletion from the primary sample and acquired an amplification in *AURKA* in the metastatic sample (**Fig. 4h**). Interestingly, we observed a high-density kataegis region in chromosome 19, within the gene *CIC*, which presented a deep deletion in both the primary and the metastatic samples (**Fig. 4i**). The proximity of such clustered mutations that are highly associated with APOBEC3 activity, hints at a role for APOBEC3 mutagenesis in contributing to some such CNAs in addition to the point mutations. We also detected evidence of one such CNA event in our APOBEC3-positive cells tested for therapy resistance. In this case, upon A3A WT overexpression, the daughter cells (dA3A^WT^-5) acquired a loss of heterozygosity (LOH) event in chromosome 13 (**Fig. 4j**). Similar chromosome 13 LOH was present in 3/6 APOBEC3-positive cells but none of the 10 APOBEC3-negative cells (*p*=0.035). When dA3A^WT^-5 cells were exposed to the CDK4/6i, abemaciclib, they selectively acquired APOBEC3-context mutations in the *RB1*. Q217* was the most frequent SNV observed in *RB1* in our clinical cohort, enriched in APOBEC3-dominant cases (**Fig. 4e**). The lone non-APOBEC3 dominant case with this mutation nonetheless exhibited 63.5% of APOBEC3 signature contribution, suggesting a causal role of APOBEC3 mutagenesis in inducing this truncating mutation. These loss-of-function mutations in *RB1* led to a complete loss of RB1 protein expression (**Fig. 4k**) and insensitivity to abemaciclib. The resistance was also confirmed by a ∼31-fold increase in IC50 value of the resistant cells compared to DMSO-treated cells (**Fig. 4l**). Overall, these cases illustrate both pre-existence of APOBEC3 mutagenesis prior to drug exposure, but also the marked accumulation and diversification over time arguing for APOBEC3 being an active mutagenic process. Taken together with the clinical observations, our data reveal that APOBEC3 mutagenesis specifically leads to APOBEC3-class alterations that drive resistance to breast cancer therapies and promote lethal outcomes.

### Targeting APOBEC3-enriched breast cancers

Our results establish a causal role of APOBEC3 mutagenesis in promoting resistance in HR+/HER2-breast cancers revealing the necessity to improve targeting of this specific class of tumors with an active APOBEC3 mutagenic processes. In the absence of reliable APOBEC3-targeting agents, we sought to determine whether we could exploit the specific biomarkers enriched in APOBEC3-dominant tumors that serve as indications for targeted therapies (*PIK3CA* mutation for PI3Kα-selective inhibitors^51^ and high TMB for anti-PD-1 monoclonal antibodies^52^). We identified 39 TMB-high, HR+/HER2-advanced breast cancer patients who received anti-PD-1 immunotherapy. Although no statistically significant difference in median PFS was observed in APOBEC3-dominant versus non-APOBEC3-dominant tumors (1^st^/2^nd^ line: 5.1 vs 3.9 months, *p*=0.6, >2^nd^ line: 1.7 vs 1.7 months, *p*=0.9, **Fig. S7**), patients receiving immunotherapy in early settings (1^st^/2^nd^ lines) displayed a numerically longer PFS than patients receiving immunotherapy in later lines. Among the earlier treated patients (*n* = 14), all patients with APOBEC3-dominant tumors showed disease control (**Fig. 5a**). We next examined patients with HR+/HER2-advanced breast cancer who received PI3Kα-selective inhibitors for treatment of metastatic disease. Although numbers were limited to effectively compare PFS based on dominant signatures (<4 lines, *n* = 21), there were several APOBEC3-dominant cases on this therapy for over 6 months (n = 5/8), with one patient responding for more than 30 months (**Fig. 5b**). Consistent with this, we found a higher rate of double *PIK3CA* mutation accumulation among APOBEC3-dominant cases compared to non-APOBEC3 cases, regardless of TMB category (7.6% vs 2.5%, *p*<0.001, **Fig. 5c**), indicating an active role for APOBEC3 mutagenesis in tumor evolution.

**Fig 5.**
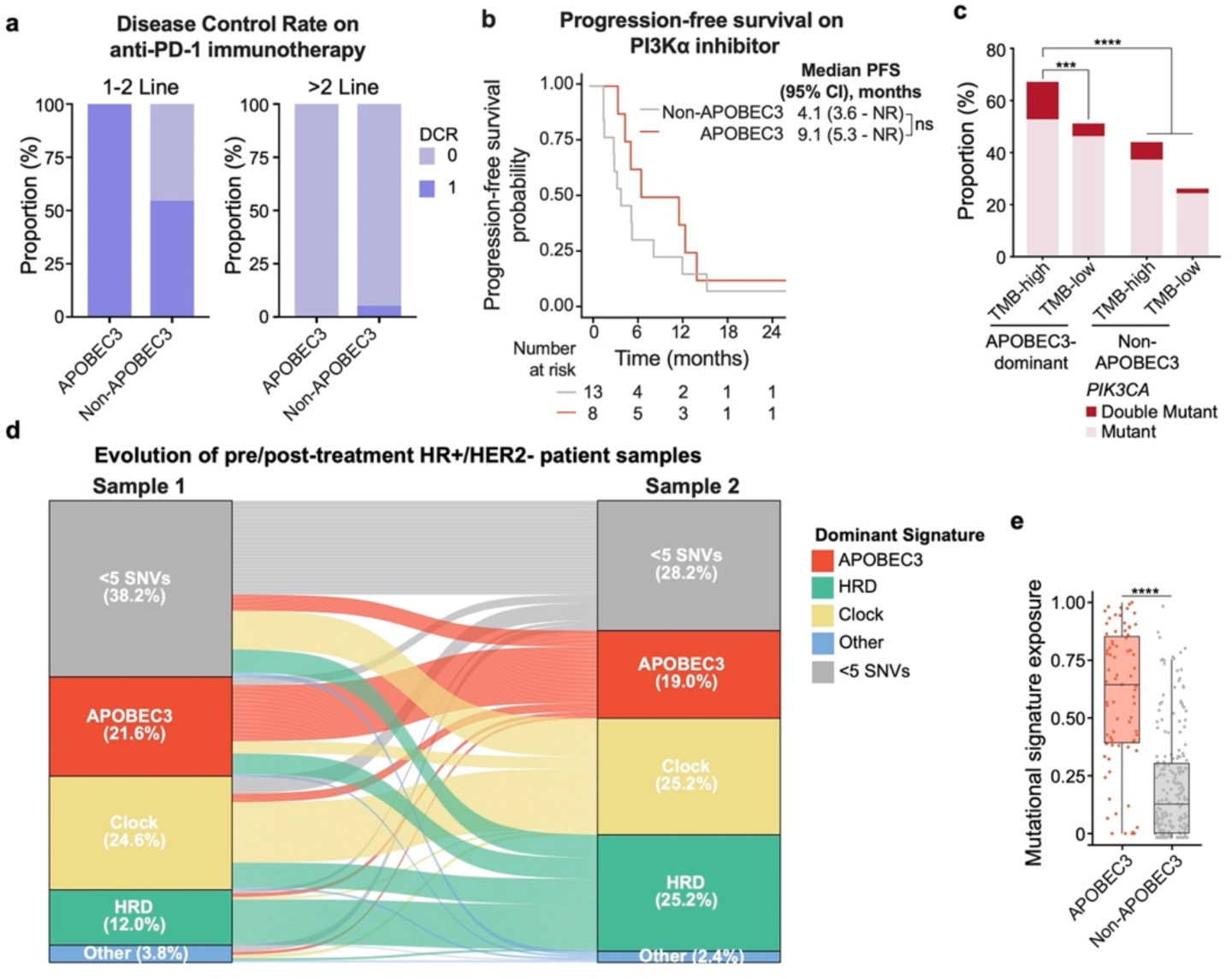
Vulnerabilities of APOBEC3-dominant breast cancers. **(a)** Barplots displaying the proportion of APOBEC3-dominant and non-APOBEC3 dominant tumors categorized based on the disease control rate (DCR) on 1-2 lines or >2 lines of anti-PD-1 immunotherapy. **(b)** Kaplan-Meier curves displaying progression-free survival probability of patients with HR+/HER2-metastatic breast cancers treated with PI3Kα inhibitor. The patients are categorized according to the dominant mutational signatures of the biopsy obtained prior to treatment start. The groups were compared using log-rank test: ns not significant. **(c)** Barplots representing proportion of *PIK3CA* mutations in APOBEC3-dominant and non-APOBEC3 dominant tumors categorized as TMB-high or TMB-low. The groups were compared using the Fisher’s exact test: *** *p*<0.001, **** *p*<0.0001. **(d)** Sankey plots representing the evolution of early/late paired HR+/HER2-patient samples categorized according to the dominant mutational signatures. **(e)** Boxplots displaying the mutational signature contribution in paired earlier samples when the later sample is APOBEC3-dominant. The groups were compared using the two-tailed Wilcoxon test: **** *p*<0.0001.

Finally, given the potential utility of earlier identification and use of targeted and combination therapies in breast cancer, we sought to establish whether APOBEC3 signatures might be evident earlier in the course of disease. To test this, we evaluated the evolution of APOBEC3 mutational signatures in the paired cohort (**Fig. 5d**). Among cancers where APOBEC3 was the dominant signature in the later timepoint, over 60% showed APOBEC3 as the dominant signature in the earlier timepoint with a median APOBEC3 signature exposure of 64% (IQR, 39-85%, **Fig. 5e**). These findings reveal a persistent role for APOBEC3 mutagenesis in most cases, rather than a *de novo* acquisition during treatment exposure. The findings thus highlight APOBEC3 signatures as early biomarkers for poor outcomes underscoring their potential for deployment in identifying tumors at risk for later development of treatment resistance.

## DISCUSSION

In this report, we demonstrate that the population of treatment refractory, poor prognosis breast cancers are marked by pervasive APOBEC3 mutagenesis. In both patient samples and laboratory models, APOBEC3 enzymes A3A and A3B drive tumor evolution and promote treatment resistance by inducing APOBEC3-context alterations in key resistance-associated genes. While APOBEC3 activity is observed to promote new, subclonal alterations that mediate treatment resistance, evidence for APOBEC3 activity is commonly detectable in pre-treatment samples, suggesting the utility of APOBEC3 signatures as biomarkers as well as revealing the potential for alternative approaches to targeting these high risk cancers.

While identification of the specific mutations that mediate therapy resistance in advanced cancer has led to the development of improved second-generation inhibitors or drug combinations, the capacity for a large subset of cancers to continually evolve and subvert second or third generation inhibitors has implied the need for better understanding of the specific processes of tumor evolution that these evolvable cancers utilize to achieve such resistance. Such understanding has the potential both to identify the cancers at risk for future development of treatment resistance, but also suggest the types of treatment approaches capable of overcoming these future resistance trajectories. In the case of breast cancer, the widespread prevalence of the APOBEC3 mutational signature among metastatic tumors suggested its relevance for tumor evolution and we therefore set out to examine its specific function in this crucial clinical context.

First, by examining a cohort of patients for which detailed treatment histories are available, we were able to establish that APOBEC3 mutagenesis is indeed associated with shorter outcomes on targeted therapies. This was true for both antiestrogen monotherapy and antiestrogen plus CDK4/6 kinase inhibitor therapy among HR+ breast cancers. Whether the same could be said for other regimens or disease subsets would likely require larger cohorts for those subsets. To evaluate causality of APOBEC3 mutagenesis in resistance, we utilized laboratory models in which APOBEC3 mutagenesis was introduced or else models in which extant APOBEC3 mutagenesis was abrogated. In both cases, APOBEC3 mutagenesis reproducibly accelerated the capacity of tumor cells to develop resistance. Moreover, APOBEC3 mutagenesis was, in some models, found to drive resistance by introducing the same types of alterations that are observed in patients that have acquired therapy resistance such as the loss-of-function mutations in *RB1* after exposure of HR+ cells to CDK4/6 inhibitors. Moreover, the clinical data point to a set of known resistance alterations (e.g., *ESR1* E380Q or *ARID1A* truncating mutations), among APOBEC3 signature positive tumors generated in the A3A or A3B context. These data together effectively establish a role for APOBEC3 activity in mediating therapy resistance, even while generating new questions such as what additional types of genomic alterations are enriched in APOBEC3-dominant tumors, are A3A and/or A3B directly responsible for mediating these alterations, or when in the course of disease are these forms of tumor evolution occurring.

The question of timing of APOBEC3 mutagenesis has been somewhat controversial as there has been evidence both for persistent and staggered events^42^. Moreover, the enrichment of APOBEC3 mutagenesis in metastatic breast cancers compared to primary breast cancers has further led to the question of whether the APOBEC3 activities are ‘acquired’ akin to the acquired mutations such as *ESR1* that are observed or simply reflect the more aggressive and evolvable cancers that recur as metastatic disease. The ability to profile a large cohort of paired primary and metastatic tumors provides some answers to this question. By analyzing paired primary and metastatic cases, we identified that 60% of APOBEC3-dominant metastatic tumors had evidence for APOBEC3 signatures in the primary. Moreover, A3A or A3B protein expression at some level was commonly observed in most APOBEC3-dominant tumors (95%), further lending support to the notion that evidence for APOBEC3 activity can be detectable at an earlier stage. Indeed, this is consistent with our finding of evidence for APOBEC3 signatures even in some pre-invasive breast lesions^31,53^. This is distinct from other cancer types where APOBEC3 activation may be initiated by targeted therapy itself^54,55^. The preexistence of APOBEC3 in breast cancer critically establishes the potential for the development of biomarkers that can detect the underlying evolvability of a breast cancer even prior to its exposure to therapy and development of the acquired mutations that mediate resistance. Such biomarkers are likely to be the first step towards enabling drug treatment strategies that overcome resistance as well as limiting the use of unnecessary therapies to those not at risk of such evolution.

In terms of the types of therapies that might be of greatest use against breast cancers harboring APOBEC3 mutagenesis, there has been limited evaluation of therapies specifically for this context. The genomics patterns of APOBEC3-dominant tumors do reveal two potentially interesting targets. First, APOBEC3-dominant tumors often harbor mutations in the helical domain of *PIK3CA* and are also enriched in double *PIK3CA* mutations. The preponderance of *PIK3CA* alterations may relate to a unique dependence on the PI3K pathway for APOBEC3-dominant cancers and suggest a subset uniquely vulnerable to PI3K-targeting therapy in the earlier stages prior to evolution to specific forms of resistance such as acquired *PTEN* or *AKT1* mutations^11,56^. Second, APOBEC3-dominant tumors often display high TMB, an indication for immune checkpoint blockade across solid tumors^30^. Indeed, a subset of patients treated under this indication were found to be APOBEC3-dominant, and among these several prolonged responses were observed in the metastatic, treatment-refractory setting. Given the potential for improved efficacy of immunotherapies in early-stage breast cancer compared to the metastatic, treatment-refractory setting, it is intriguing to speculate whether presence of APOBEC3 activity represents a potential indication for such therapies in a subset of early-stage tumors. This potential is made all the more poignant by the recent reporting of an improved rate of pathologic complete response for the combination of chemotherapy with pembrolizumab for a subset of ER-positive early-stage breast cancer^57^. However, even beyond these possibilities, the use of many existing therapies (CDK4/6 inhibitors, antibody drug conjugates, etc.) may potentially benefit from a mutational signature-based biomarker, revealing additional evidence of a cancer with the potential for further evolution. The study presented here has several notable limitations that will be important to address with future research. First, the large clinic-genomic cohort was evaluated by MSK-IMPACT targeted panel with materially reduced information than WGS/WES analysis. Although we have taken several steps to ensure the fidelity and reproducibility of our findings on APOBEC3 mutagenesis with the publicly-availably datasets, the use of WGS will likely enable further understanding of the scope and significance of APOBEC3 mutagenesis in the clinic. Another limitation is the preponderance of ER-positive tumors in our clinical cohort, reducing our ability to ascertain how APOBEC3 mutagenesis contributes to disease progression in the other breast cancer subtypes. Our HER2+ preclinical models do hint at a role for APOBEC3 mutagenesis in this subtype, but a comprehensive analysis with a larger sample size is needed to determine whether and how APOBEC3 activity mediates resistance to HER2-targeted therapies in patients.

In closing, we reveal that APOBEC3 mutagenesis represents a highly prevalent driver of genomic instability in breast cancer that specifically contributes to the development of resistance to endocrine and targeted therapies. Our results further demonstrate that the presence of APOBEC3 activity can be detected before exposure to therapies in breast cancer, and may therefore, represent a valuable biomarker and therapeutic target in this disease.

## METHODS

### Study cohort

A total of 5,831 breast cancer samples, which underwent prospective genomic profiling by the Memorial Sloan Kettering-Integrated Mutation Profiling of Actionable Cancer Targets (MSK-IMPACT) targeted sequencing panel^4,58,59^ from January 2014 and December 2021, were retrieved. After removing samples with a sequencing-estimated tumor purity <20% (see below), a total of 3,880 breast cancer samples from 3,117 patients were included. The study was approved by the Memorial Sloan Kettering Cancer Center (MSKCC) Institutional Review Board (12-245) and all patients provided written informed consent for tumor sequencing and review of medical records for demographic, clinical and pathology information. Demographics, pathologic and detailed clinical information was collected until date of data freeze (June 2022).

Histological subtypes, tumor stage at diagnosis, tumor grade of primary breast cancer, and receptor status were determined as described in *Razavi et al*^4^. Briefly, breast cancer histological subtypes were classified as either invasive ductal carcinoma (IDC), invasive lobular carcinoma (ILC), or mixed/other histologic types. Tumor grade was defined based on the Nottingham combined histologic grade of the primary breast cancer. The primary tumors with total tumor score of 3-5 were classified as G1 (well differentiated); 6-7: as G2 (moderately differentiated), and 8-9: as G3 (poorly differentiated). Patients were classified into breast cancer subtypes based on ER and PR IHC results and the HER2 IHC and/or FISH results rendered at the time of diagnosis in accordance with the American Society of Clinical Oncology (ASCO) and College of American Pathology (CAP) guidelines^60,61^.

Each patient in our cohort was assigned a singular receptor status. Recognizing potential intertumoral heterogeneity, we sought a unified definition as follows: i) in cases where any metastatic biopsy was sequenced, receptor status was defined by treating clinician interpretation and assigned first-line treatment; ii) in cases where only a primary tumor is sequenced, receptor status was defined by receptor status of the sequenced primary. Employing these definitions, our cohort consisted of 2,133 patients (68.4%) with HR+/HER2-tumors, 276 (8.9%) with HR+/HER2+ tumors, 149 (4.8%) with HR-/HER2+ tumors, and 559 patients (17.9%) with TNBC.

### MSK-IMPACT targeted sequencing analysis

Breast cancer samples underwent tumor-normal sequencing by the FDA-cleared MSK-IMPACT assay, which targets between 341 (2014) and 505 (2021) cancer-related genes. Genomic data extracted from MSK-IMPACT included somatic single nucleotide variants, copy number alterations (CNAs), structural variants, and additional genomic metrics as tumor mutational burden (TMB, Mut/Mb) and fraction of genome altered (FGA; i.e., the percentage of the genome affected by CNAs^58,59^). Somatic mutations were classified as pathogenic, likely pathogenic or predicted oncogenic as defined by OncoKB annotation^62^. We used FACETS^63^ to define the allele-specific gene amplifications and homozygous deletions, tumor purity, and ploidy, as previously described^64^. Whole genome doubling (WGD) status was inferred from MSK-IMPACT sequencing data, as previously described^64^. Briefly, tumor samples were considered to have undergone WGD if the fraction of major allele >1 was >50%.

### Validation of *SigMA* Performance on MSK-IMPACT and cell line WGS data

While *SigMA* was originally designed to identify tumors positive for HRD, it has also proven effective in accurately classifying other types of mutational signatures, particularly excelling with APOBEC3 signatures^23^. SBS2 and SBS13 exhibit distinct trinucleotide context spectra, strongly differing from flat signatures like SBS3 and SBS5, or other specific signatures such as POLE and SMOKING, even with a minimal number of mutations for analysis. We thus aimed to independently assess *SigMA*’s ability to detect APOBEC3 signatures in breast cancer samples across three distinct datasets: TCGA breast cancers (primary, WES), *Bertucci et al.*^18^ (metastatic, WES), and *Nik-Zainal et al.*^26^ (primary, WGS). To validate *SigMA*, we adopted the same simulation approach as its developers, which involves MSK-IMPACT panel simulation.

This method is based on the concept that a high-quality signature evaluation requires a substantial mutation count, achievable only through WES and WGS. We initially computed signatures using real WES and WGS data, considering these results as the ground truth. Subsequently, we simulated MSK-IMPACT samples by reducing the mutations in each WES and WGS sample to only those within genomic regions covered by the MSK-IMPACT panel (version impact 468). *SigMA* was then applied to these simulations to determine exposure and the dominant signature. Finally, we compared these outcomes with the established ground truth to obtain performance metrics, focusing on the ability to predict APOBEC3-dominant samples. Only samples with ≥SNVs in the simulated panel were considered for performance metric analysis.

### Whole genome sequencing (WGS) analysis

To validate findings on MSK-IMPACT and provide further information regarding the evolution of APOBEC3 mutagenesis over time, WGS was carried out on five pairs of selected primary and metastatic breast cancer patient samples by the MSKCC’s Integrated Genomics Operations using validated protocols^23,25^. Briefly, microdissected tumor and germline DNA were subjected to WGS on HiSeq2000 (Illumina). The median sequencing coverage depth of 102x (range, 88x-132x) for tumor and 44x (range, 36x-58x) for normal samples. In addition, genomic DNA was extracted from 29 cell lines using the E.Z.N.A Tissue DNA Extraction Systems (Omega Bio-Tek 101319-018), and subjected to WGS, with a median sequencing coverage depth of 63x (for T47D lines) or 33x (for BT-474 lines) (range, 28x-103x).

For samples and cell lines subjected to WGS, the data were processed through a validated bioinformatics pipeline^23,25^. Initially, sequence reads were aligned to the human reference genome GRCh37 utilizing the Burrows-Wheeler Aligner (BWA, v0.7.15)^65^. SNVs in both WGS and MSK-IMPACT analyses were identified using MuTect (v1.0)^66^. Insertions and deletions (indels) were detected by employing a suite of tools: Strelka (v2.0.15)^67^, VarScan2 (v2.3.7)^68^, Platypus (v0.8.1)^69^, Lancet (v1.0.0)^70^, and Scalpel (v0.5.3)^71^. CNAs and LOH assessments were conducted using FACETS^63^. Mutations in tumor suppressor genes deemed deleterious/loss-of-function, or those targeting a known mutational hotspot in oncogenes, were classified as pathogenic. Hotspot-targeting mutations were annotated with reference to cancerhotspots.org^72^.

Structural variants were identified using Manta^73^, SvABA^74^ and Gridss^75^ from WGS data. Structural variants identified by at least two of the three callers were retained and utilized for subsequent analyses. Setup and call procedures are described in detail in their respective code repositories: for Manta – https://github.com/ipstone/modules/blob/master/sv_callers/mantaTN.mk, and for SvABA: https://github.com/ipstone/modules/blob/master/sv_callers/svabaTN.mk. These processed structural variant calls, along with additional genomic data (SNVs, indels, CNAs), were integrated to create circos plots via the signature.tools.lib R package^76^ (code repository: https://github.com/Nik-Zainal-Group/signature.tools.lib). For all patients and cell lines with more than 1 sample, all unique variants from any samples in a given patient or cell line were genotyped in all other samples from the same patient or cell line using Waltz (https://github.com/mskcc/Waltz).

### Mutational signature analysis

We have applied various tools to compute mutational signatures across different types of data. For WES data, DeconstructSigs^29^, MutationalPatterns^28^, and SigProfiler^27^ were employed. For WGS data, we opted for Signal^77^, and for the MSK-IMPACT panel, *SigMA* was used. The DeconstructSigs method is designed to identify the most accurate linear combination of predefined signatures that reconstructs a tumor sample’s mutational profile. This method employs a multiple linear regression model. DeconstructSigs, an R package extension, leverages the Bioconductor library at CRAN (https://cran.r-project.org/). MutationalPatterns operates as a non-negative least squares (NNLS) optimization algorithm. The NNLS problem is extensively studied, and MutationalPatterns utilizes an R-based active set method from the pracma package for its ‘fit_to_signatures’ function, available at CRAN. We ran MutationalPatterns with two different settings, ‘regular’ and ‘strict’. The ‘strict’ method suffers less from overfitting but can suffer from more signature misattribution. SigProfilerSingleSample attributes a set of known mutational signatures to an individual sample, determining the activity of each signature and the probability of each causing specific mutation types. It integrates SigProfilerMatrixGenerator and SigProfilerPlotting for its functionality. We also utilized SigProfilerSimulator and SigProfilerClusters to assess the clustered mutations. Signal is considered one of the best state-of-the-art tool for WGS, offering a comprehensive workflow for mutational signature analysis. Kataegis was detected using the Signal web portal (https://signal.mutationalsignatures.com).

*SigMA* can process samples with at least 5 somatic SNVs, making it particularly suitable for MSK-IMPACT samples. It encompasses five steps: discovery of mutational signatures in WGS data using NMF; clustering to determine tumor subtypes; simulation of cancer gene panels and exomes; calculation of likelihood, cosine similarity, and signature exposure; and training of gradient boosting classifiers for a final score. *SigMA* score thresholds are established based on simulated data, considering tumor type and sequencing platform.

The dominant mutational signature in each MSK-IMPACT sample, i.e., the primary mutational process occurring in a cancer genome, was assessed using *SigMA*. We translated mutational exposures into percentages for cross-sample comparisons. For all other tools used in WGS and WES data, the dominant signature was defined as the mutation process with the highest percentage of exposure. This included groupings as Clock (SBS1+SBS5), APOBEC3 (SBS2+SBS13), HRD (SBS3+SBS8), and considering Other signatures independently (e.g., SBS17, SBS18, etc.). The process with the maximum exposure was considered dominant. To determine if a mutation was APOBEC3-context, we identified characteristic peaks from cosmic signatures related to SBS2 (C>T mutations in the contexts TCA, TCC, TCG, TCT) and SBS13 (C>G mutations in the contexts TCA, TCC, TCG, TCT; C>A mutations in the contexts TCA, TCC, TCG, TCT), as described previously^23^.

### Treatment outcome analysis

All patients included in the treatment outcome analyses were treated at MSKCC. The exact regimen, dates of start and stop therapy, as well as date of progression was annotated via expert review. Progression events were defined as i) radiographic or clinical event prompting change in systemic therapy or recommendation for locally targeted radiation therapy, ii) documented clinician impression detailing progression, after which there was documented patient or MD preference to continue same therapy, as previously described^4^.

We determined the association between dominant mutational signature and PFS with disease progression on therapy with endocrine therapy +/- CDK4/6 inhibitors or patient death. Endocrine therapies, either as monotherapy or as partner of CDK4/6 inhibitors were categorized as following: aromatase inhibitor vs selective estrogen receptor degrader. Log-rank test was used to compare the survival distributions among two or more groups. Both univariate and multivariate Cox proportional hazard models (stratified by endocrine therapy partner, and treatment line, where available) were applied. For patients with multiple lines of therapy from the same class of treatment, only the first treatment line from that class that was started after the MSK-IMPACT biopsy was included in the analysis.

### Permutation analysis for APOBEC3 enrichment

To assess the correlation between APOBEC3 enrichment and the occurrence of SNVs within a gene, we adapted a method previously described^78^ for our dataset. The aim was to detect potential correlations between mutations in specific genes and APOBEC3 enrichment. To mitigate the impact of high TMB in APOBEC3-dominant samples, high frequency of mutations in longer genes, or the presence of high frequency genetic alterations in breast cancer, we applied a permutation-based method aimed at standardizing the overall mutation count for each sample and gene. This process randomized the gene × sample binary mutation matrix while preserving the mutation counts for each gene and sample, following the approach described by *Strona et al*^79^. In this matrix, rows represent samples and columns represent genes, with ‘1’ indicating a mutation and ‘0’ indicating a wild-type non-mutated gene. We conducted an APOBEC3 enrichment analysis across all genes using a Wilcoxon rank-sum *p* value to compare APOBEC3 exposure in mutant versus wild-type samples for each gene. To correct for multiple testing, *p* values were adjusted for the false discovery rate using the Benjamini-Hochberg procedure. For the original dataset and each of 10,000 permutations, we calculated the Pvalue_observed and Pvalue_Random, respectively (for a total of 10,000 Pvalue_Random). The final *p* value was calculated as the number of permutation iterations where Pvalue_Random ≤ Pvalue_observed, divided by the total number of permutations (10,000), ensuring a fair assessment of gene-mutation frequencies across samples with high mutation burdens. For the permutation analysis, we utilized the R package EcoSimR. The permutation test was carried out under various conditions, separately for HR+/HER2- and for TNBC in both primary and metastatic settings, using pathogenic and likely-pathogenic SNPs by OncoKB annotation. For each data group, we filtered for genes mutated in at least 1% of the subset obtained and for samples with at least one mutation in one of these genes. This filtering was necessary to ensure the EcoSimR package performed correctly during the randomization process. To compute *p* values, odds ratio, false discovery rates and other statistics we used different functions from the Python packages *scipy* and *statsmodels*, while for visualization we used *matplotlib*.

### Genomic analysis of patients with multiple samples collected over time

To investigate the evolution of mutational processes over time, we subset the initial cohort of 3,800 breast cancers from MSK-IMPACT identifying patients with HR+/HER2-subtype and multiple tumor samples collected over time and various treatments (patients with 2 samples, n = 449; patients with ≥3 sample, n = 43). Tumor sample pairs with complete somatic mismatches on the levels of single nucleotide variants, indels, and/or CNAs were excluded.

To assess whether a somatic mutation was truly acquired over time and exposure to therapy, somatic mutations identified in samples collected afterward were interrogated in the matched respective primary tumor/first metastatic biopsy in a matched tumor-informed manner (genotyping) using Waltz (https://github.com/mskcc/Waltz), which required at least 2 duplex consensus reads, comprising both strands of DNA, to call a somatic SNV at a site known to be altered in the matched tumor sample from a given patient, as previously described^80^.

### Immunohistochemical analysis

Immunohistochemical analyses were conducted on a Leica Bond III automated stainer platform (Leica, Deer Park, IL). Four-micron thick formalin-fixed paraffin-embedded (FFPE) tissue sections were subjected to heat-based antigen retrieval for 30 minutes using a high pH buffer solution (Bond Epitope Retrieval Solution 2; Leica, AR9640). Subsequently, they were incubated with the A3A-13 (LQR-2-13 (UMN13)) or the A3B (5210-87-13) primary antibodies at a 1:2,500 and 1:200 dilution, respectively for 30 minutes. A polymer detection system (Bond Polymer Refine Detection; Leica, DS9800) was used as secondary reagent. Extent (% of tumor cells) and intensity (weak, moderate, strong) of A3A and A3B expression was evaluated was evaluated by two pathologists (FP and JSR-F).

### Cell lines

The following cell lines were used: T47D (ATCC HTB-133), MCF7 (ATCC HTB-22), BT-474 WT and *A3A* KO, and MDA-MB-453 WT and *A3A* KO (gift from John Maciejowski) and HEK293T (gift from Ping Chi). T47D cells were cultured in RPMI; MCF7, BT-474 and MDA-MB-453 were cultured in DMEM/F12; HEK293T were cultured in DMEM media. All media were supplemented with 10% fetal bovine serum (Corning 35-010-CV), 2 mM L-glutamine, 100 U/mL penicillin, and 100 µg/mL streptomycin (Gemini Bio 400-109). Tetracycline-free fetal bovine serum (Takara Bio 631367) was used for experiments using the doxycycline-system. All cells were maintained in a humidified incubator with 5% CO_2_ at 37°C. All cell lines were routinely tested negative for mycoplasma contamination.

### Cloning and lentiviral transduction

The A3A^WT^, A3A^E72Q^, A3B^WT^ and A3B^E255Q^ sequences (cDNA plasmids gift from John Maciejowski) were cloned into pDONR221 (Invitrogen 12536017) using the Gateway BP Clonase II Enzyme mix (Invitrogen 11789020). Expression vectors were generated by cloning into pInducer20 (gift from Stephen Elledge, Addgene plasmid 4401^81^) using the Gateway LR Clonase II enzyme mix (Invitrogen 11791020). Lentiviral particles were prepared by transfecting 293T cells with expression clones along with the lentiviral envelope and packaging plasmids using X-tremeGENE HP DNA transfection reagent (Sigma-Aldrich 6366546001). The media was refreshed after 24 h. After 48 h, the supernatant was collected and filtered through 0.45 µm filters. T47D and MCF7 cells were transduced with the lentiviral particles and stably expressing cells were generated after selection using G418 (InvivoGen ant-gn-1) for two weeks. Cells were treated with 0.1 µg/mL doxycycline (Sigma-Aldrich, D9891) to induce overexpression.

### Generation of resistant cells

T47D dA3A^WT^-5 and dA3A^E72Q^-5 cells were treated with DMSO or 500 nM abemaciclib for 14 days, after which the drug concentration was increased to 1 µM. Surviving dA3A^WT^-5 cells were tested for resistance after 2 months of continuous culture in 1 µM abemaciclib. dA3A^E72Q^-5 cells did not grow out even after 5 months under selection. Additionally, resistant cells were expanded after selection from the long-term growth assays.

### Antibodies and reagents

The following primary antibodies were obtained from Cell Signaling Technology and used for immunoblotting at a dilution of 1:1000: anti-HA (C29F4), anti-p-Rb S780 (D59B7), anti-p-Rb S807/811 (D20B12), anti-Rb (4H1), anti-E2F1 (3742), anti-Cyclin E2 (4132S), anti-Cyclin A2 (BF683), and anti-Vinculin (E1E9V). The secondary antibodies used were anti-Rabbit IgG, HRP-linked (1:3000, Cell Signaling Technology 7074), anti-Rabbit IgG IRDye 680 RD (1:10000, LI-COR Biosciences 926-68071), and anti-Mouse IgG IRDye 800 RD (1:10000, LI-COR Biosciences 926-32210). The following drugs were purchased from Selleck Chemicals: fulvestrant (S1191), abemaciclib (S5716), palbociclib (S1579), lapatinib (S2111), neratinib (S2150), and MK2206 (S1078), and dissolved in DMSO. Trastuzumab deruxtecan (DS-8201a, Daiichi Sankyo) was dissolved in saline.

### Immunoblotting

Cells were lysed on ice using RIPA lysis buffer (Thermo Scientific 89901) supplemented with protease and phosphatase inhibitor (Thermo Scientific 78444). Lysates were cleared using centrifugation and total protein concentration was measured with BCA protein assay (Thermo Scientific 23225). 30-50 µg total protein was separated on 4-12% Bis-Tris protein gels (Invitrogen NuPAGE) and transferred onto PVDF or nitrocellulose membranes. Blots were blocked with Intercept (TBS) Blocking Buffer (LI-COR Biosciences 927-60001) or 5% milk, and incubated with primary antibodies overnight at 4°C. After incubation with secondary antibodies, the blots were scanned by Odyssey CLx Imaging System (LI-COR Biosciences) or developed using the Western Lightning Plus-ECL (PerkinElmer NEL104001EA).

### DNA deaminase assay

*In vitro* deamination assay was performed as described previously^82^. Briefly, 50 µg total cell lysates were incubated with RNase A (1.75 U), ssDNA substrate (4 pmol), UDG buffer and UDG (1.25 U) in HED buffer (25 mM Hepes pH 7.8, 5 mM EDTA, 10% glycerol, 1 mM DTT freshly supplemented with protease and phosphatase inhibitor) for 2 h at 37°C. 100 mM NaOH was then added and heated at 95°C for 10 min to cleave the DNA at abasic sites. The sample was then heated with Novex Hi-Density TBE sample buffer (Invitrogen LC6678) at 95°C for 5 min, cooled down on ice and run on a 15% TBE-urea PAGE gel. Separated DNA fragments were imaged on an ImageQuant 800 (Amersham). The ssDNA oligo substrates were 5’-(6-FAM)-GCAAGCTGTTCAGCTTGCTGA for A3A^83^ and 5’-ATTATTATTATTCAAATGGAT-TTATTTATTTATTTATTTATTT-fluorescein for A3B.

### *In vitro* growth assays

For cell viability measurement, 500-2,000 cells/well were plated in triplicates in 96 well plates. After overnight incubation for attachment, cells were treated with drugs (day 0). Media was replaced once every week and doxycycline was topped up twice every week for overexpression cells. Viability was measured using the redox-sensitive dye Resazurin (R&D Systems AR002; 25 µL per well incubated at 37°C for 4 h and read at 544 nm excitation and 590 nm emission using SpectraMax M5 (Molecular Devices)) or brightfield imaging of attached cells using Incucyte S3 (Sartorius). Half maximal inhibitory concentrations (IC_50_) were calculated by nonlinear regression of inhibitor concentrations with response in GraphPad Prism 9. For colony forming assays, 2 x 10^5^ MDA-MB-453 cells were plated in triplicates in 6 well plates. The cells were treated with drugs after 24 h, and their media was replaced with fresh drugs twice every week. The cells were fixed with 100% methanol when confluent, washed with water, stained with 0.5% crystal violet (Sigma-Aldrich C0775) in 25% methanol, washed with water, dried, and scanned using the AxioObserver 7 inverted microscope (Zeiss).

### Statistical analyses

Statistical analyses were conducted using R (version 3.1.2) and GraphPad Prism v9.4.1. Summary statistics were used to describe the study population. Fisher’s exact test or Pearson’s chi-squared test were used to compare categorical variables, whenever appropriate. Mann–Whitney U or Wilcoxon rank sum, or two-way ANOVA tests were used to compare continuous variables. Log-rank test was used to compare the survival distributions between groups. Comparisons of frequencies of genes altered by somatic SNVs and CNAs as well as for site-specific gene alterations were performed using the Fisher’s exact test and logistic regression. Multiple testing correction using the Benjamini–Hochberg method was applied to control for the false discovery rate (FDR) whenever appropriate. Pearson’s coefficient *R* was computed using the python package scipy.stats. All *p* values were two-tailed, and 95% confidence intervals were adopted for all analyses.

### Data availability

The MSK-IMPACT sequencing dataset will be available through the cBioPortal for Cancer Genomics at www.cBioPortal.org at the time of publication. WGS data are available upon request from the corresponding author. WGS data from cell lines will be deposited and made available before publication. Data for breast cancers from TCGA were downloaded as the harmonized MC3 public MAF from https://gdc.cancer.gov/about-data/publications/mc3-2017, from Nik-Zainal et al. were download from the ICGC data portal (https://dcc.icgc.org), and data from Bertucci et al. were provided by Dr. Fabrice André. Source data including comparison of signature assessment from different signature calling tools, quantification of mutational signatures and clustered mutations from WGS of cell lines and patient samples, and permutation tests for gene-enrichment analyses are provided as supplementary material.

### Code availability

All software used in this manuscript are available online.

## ACKNOWLEDGEMENTS

SC acknowledges grant support from the Breast Cancer Research Foundation, NCI SPORE grant P50 CA247749, NCI Cancer Center Support Grant P30 CA008748, the Naddisy Foundation, and the Cancer Couch Foundation. Research reported in this publication was further supported in part by a Cancer Center Support Grant of the NIH/NCI (Grant No. P30CA008748). We gratefully acknowledge the members of the Molecular Diagnostics Service in the Department of Pathology, the Integrated Genomics Operation and Bioinformatics Core, and the Marie-Josée and Henry R. Kravis Center for Molecular Oncology. BW is funded in part by Breast Cancer Research Foundation, Cycle for Survival, and NIH/NCI P50 CA247749 01 grants. JSR-F was funded in part by the Breast Cancer Research Foundation, by a Susan G Komen Leadership grant, and by the NIH/NCI P50 CA247749 01 grant. Cancer studies in the Harris lab are supported by NCI P50-CA247749, NCI P01-CA234228, and a Recruitment of Established Investigators Award from the Cancer Prevention and Research Institute of Texas (RR220053). RSH is an Investigator of the Howard Hughes Medical Institute, a CPRIT Scholar, and the Ewing Halsell President’s Council Distinguished Chair at University of Texas Health San Antonio. FP is partially funded by an NIH/NCI P50 CA24779 grant and by a Starr Cancer Consortium grant.

## AUTHOR INFORMATION

### Contributions

AvG, AnG, RSH, JSR-F, AM and SC designed the study. AvG, AM and SC wrote the paper. AvG performed data curation and analyses for all experiments with the cell line models. AnG conducted the validation of *SigMA*, APOBEC3 context enrichment and permutation test analyses. PS processed all MSK-IMPACT and patient and cell line WGS data. PS, DNB, YZ, JP, JB-H and XP performed computational analysis of all sequencing data. Mutational signatures and clustered mutations analyses were conducted by AnG for the clinical and patient WGS data, and AvG for cell line WGS data. Treatment outcome analyses were performed by AM and AS. AM performed the remaining analyses on the MSK-IMPACT cohort. MAC, BS and RSH provided the antibodies for IHC, DF and AAJ performed the staining, and FP, EMS and JSR-F imaged and analyzed all samples. PR, SNP, NR, BW, GC, and ML were involved in experimental design, data interpretation or review of the manuscript. All authors approved the final manuscript.

### Materials and correspondence

Correspondence and request for materials should be addressed to Antonio Marra or Sarat Chandarlapaty.

## SUPPLEMENTARY INFORMATION

Fig S1 – Evaluation of SigMA as a tool to assess dominant APOBEC3 mutational signature

Fig S2 – Characterization of APOBEC3 mutational signatures in breast cancers

Fig S3 – A3A and A3B induce APOBEC3 mutagenesis

Fig S4 – APOBEC3 activity drives therapeutic resistance in breast cancers

Fig S5 – Gene enrichment in breast cancer samples

Fig S6 – Site-specific enrichment in HR+/HER2-breast cancer samples

Fig S7 – Outcome of HR+/HER2-breast cancers to anti-PD-1 immunotherapy

Supplementary Table 1 – Clinical and pathological characteristics of the study cohort

Supplementary Table 2 – Clinical and pathological characteristics of the study cohort by dominant signature

Supplementary Table 3 – Comparison of dominant mutational signatures assessed using different signature calling tools

Supplementary Table 4 – Signatures and clustered mutations from WGS of cell line models Supplementary

Supplementary Table 5 – Signatures and regions of kataegis from WGS of patient samples Supplementary

Supplementary Table 6 – Source data file for gene-enrichment based on dominant signatures

