## Supplementary Data for "APOBEC3 mutagenesis drives therapy resistance in breast cancer"

**SUPPLEMENTARY INFORMATION**

**Fig S1** – Evaluation of SigMA as a tool to assess dominant APOBEC3 mutational signature

**Fig S2** – Characterization of APOBEC3 mutational signatures in breast cancers

**Fig S3** – A3A and A3B induce APOBEC3 mutagenesis

**Fig S4** – APOBEC3 activity drives therapeutic resistance in breast cancers

**Fig S5** – Gene enrichment in breast cancer samples

**Fig S6** – Site-specific enrichment in HR+/HER2- breast cancer samples

**Fig S7** – Outcome of HR+/HER2- breast cancers to anti-PD-1 immunotherapy

**Supplementary Table 1** – Clinical and pathological characteristics of the study cohort

**Supplementary Table 2** – Clinical and pathological characteristics of the study cohort by dominant signature

**Supplementary Table 3** – Comparison of dominant mutational signatures assessed using different signature calling tools

**Supplementary Table 4** – Signatures and clustered mutations from WGS of cell line models

**Supplementary Table 5** – Signatures and regions of kataegis from WGS of patient samples

**Supplementary Table 6** – Source data file for gene-enrichment based on dominant signatures

### SUPPLEMENTARY FIGURES

#### Fig S1

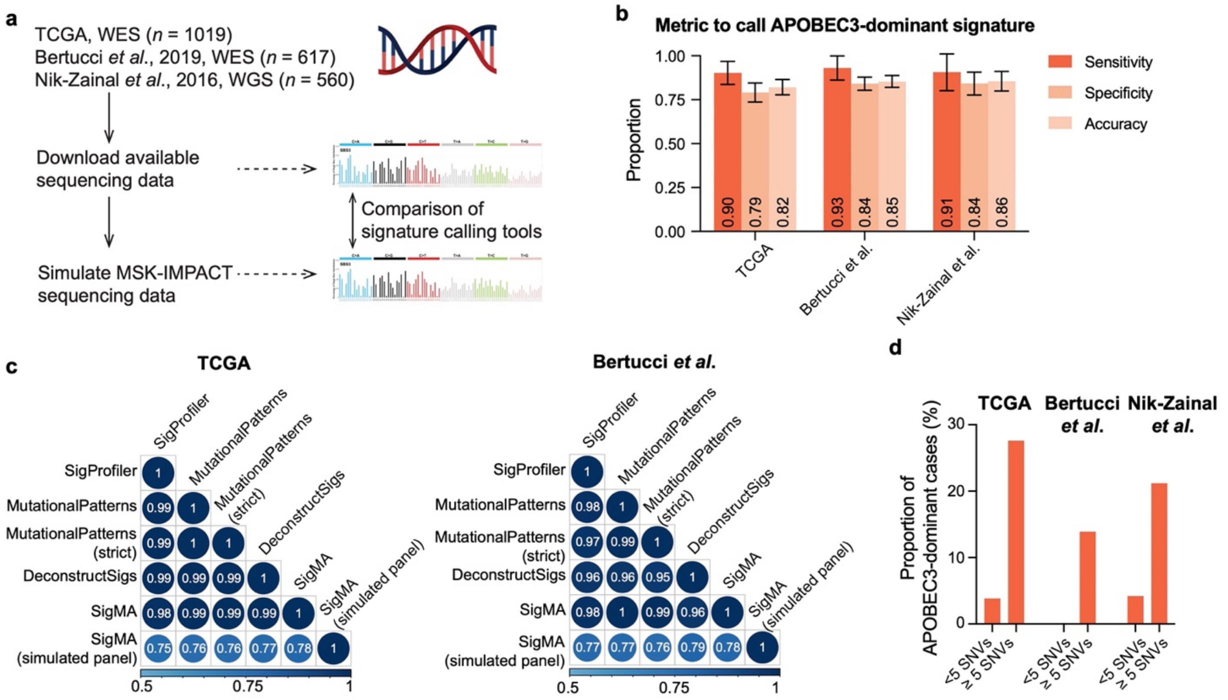


**Fig S1. Evaluation of SigMA as a tool to assess dominant APOBEC3 mutational signature. (a)** Schematic of the mutational signature analysis pipeline from publicly available datasets. **(b)** Barplots displaying the sensitivity, specificity, and accuracy of *SigMA* to call APOBEC3 as a dominant signature in the indicated datasets. **(c)** Concordance between the highlighted signature calling tools and *SigMA* in computing mutational signatures from the TCGA and Bertucci *et al.* datasets. **(d)** Barplots representing the proportion of APOBEC3-dominant samples in the indicated datasets categorized as <5 SNVs or ≥5 SNVs.

**Fig S2**

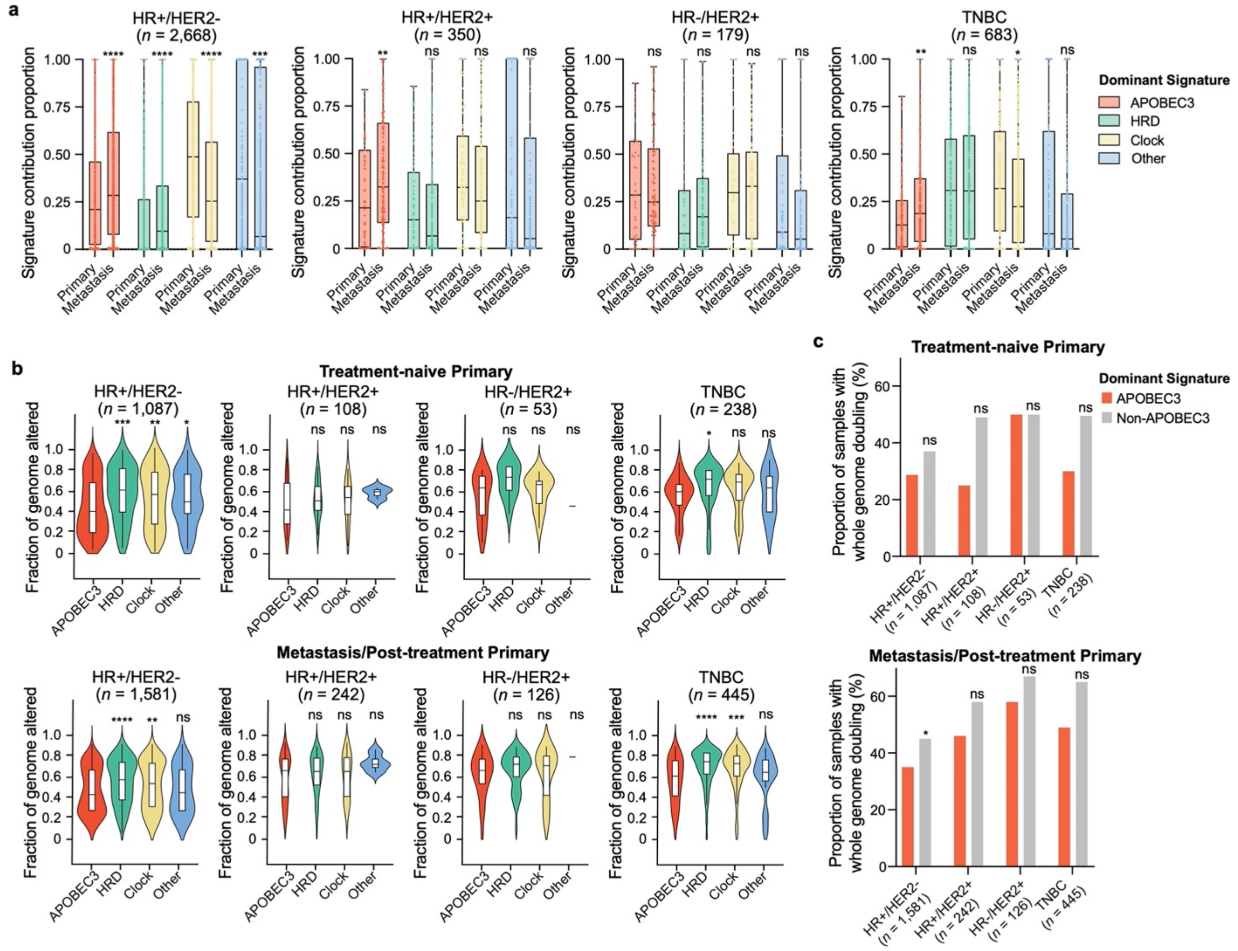


**Fig S2. Characterization of APOBEC3 mutational signatures in breast cancers. (a)** Box plots showing the contribution of APOBEC3, HRD, Clock and Other mutational signatures in primary and metastasis samples categorized according to the receptor status. The primary and metastasis groups were compared using two-tailed Wilcoxon test: ns not significant, * $p<0.05$, ** $p<0.01$, *** $p<0.001$, **** $p<0.0001$. **(b)** Violin plots displaying the fraction of genome altered in primary (top panel) and metastatic (bottom panel) patient samples categorized according to the dominant mutational signature and receptor status. The groups were compared using two-tailed Wilcoxon test: ns not significant, * $p<0.05$, ** $p<0.01$, *** $p<0.001$, **** $p<0.0001$. **(c)** Barplots representing the proportion of primary (top panel) and metastatic (bottom panel) patient samples with WGD categorized according to the receptor status. The groups were compared using two-tailed Wilcoxon test: ns not significant, * $p<0.05$.

**Fig S3**

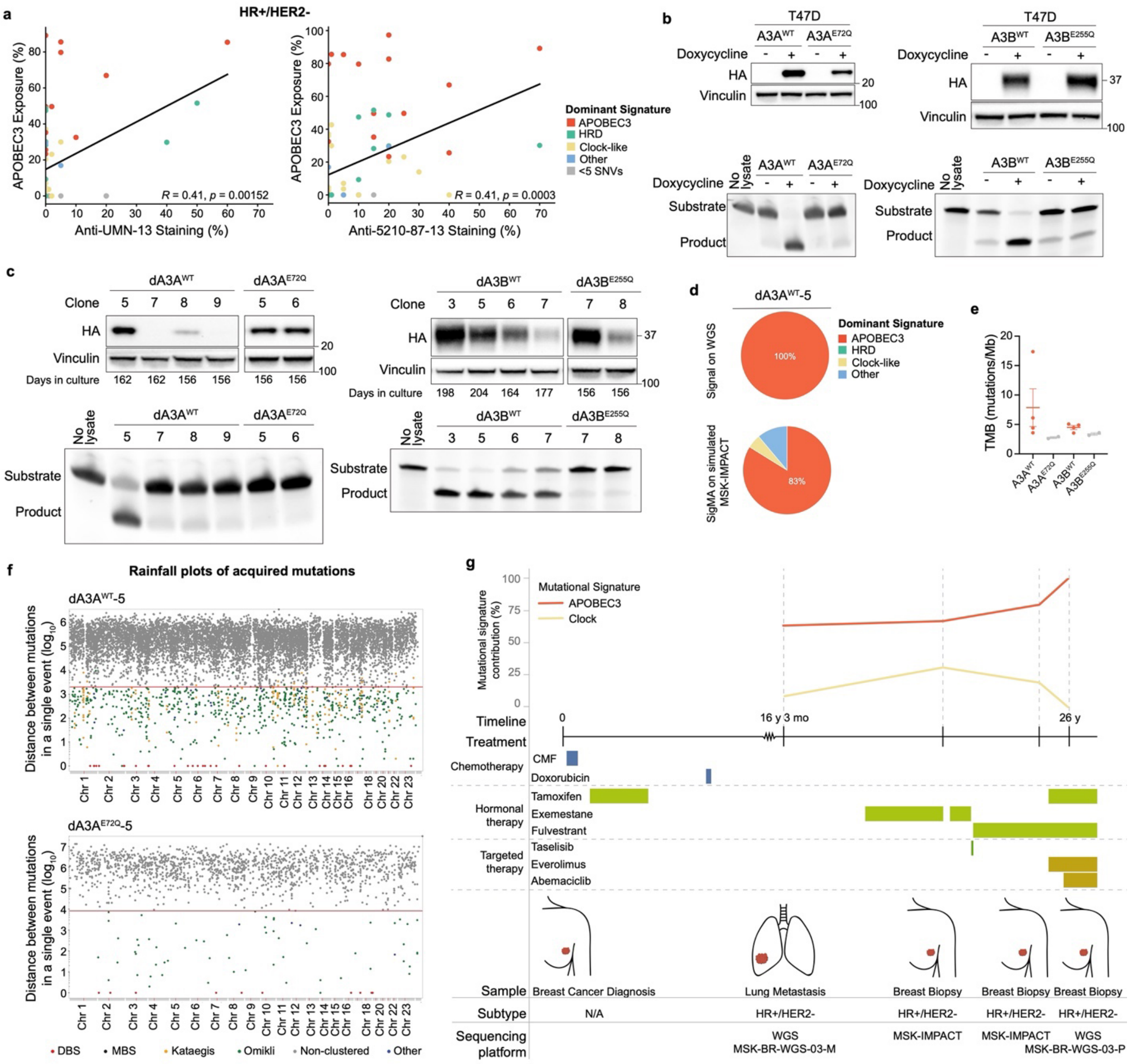


**Fig S3. A3A and A3B induce APOBEC3 mutagenesis. (a)** Scatter plot displaying the correlation between APOBEC3 signature contribution and IHC staining (in percentage of positive cells) of anti-UMN-13 (left panel) and anti-5210-87-13 (right panel) antibodies in HR+/HER2- breast cancer samples categorized according to the dominant mutational signatures. **(b)** Immunoblots of T47D cells stably transduced with doxycycline-inducible HA-tagged $A3A^{WT}$, $A3A^{E72Q}$, $A3B^{WT}$ or $A3B^{E255Q}$ (top panel). Vinculin was used as a loading control. ssDNA deaminase activity of T47D $A3A^{WT}$, $A3A^{E72Q}$, $A3B^{WT}$ or $A3B^{E255Q}$ cells (lower panel). The cells were treated with or without 0.1 µg/mL doxycycline for 48 h before harvesting. **(c)** Western blot (top panel) and ssDNA deaminase activity (lower panel) of the indicated T47D daughter clones of $A3A^{WT}$, $A3A^{E72Q}$, $A3B^{WT}$ or $A3B^{E255Q}$ cells. Vinculin was used as a loading control. The cells were treated with 0.1 µg/mL doxycycline for the indicated number of days. **(d)** Pie charts representing the contribution of mutational signatures in sample $dA3A^{WT}$-5 calculated from the WGS using Signal or the simulated MSK-IMPACT panel using *SigMA*. **(e)** TMB in the indicated samples. **(f)** Rainfall plots displaying non-clustered (grey) and clustered (colored) mutations in samples $dA3A^{WT}$-5 and $dA3A^{E72Q}$-5. **(g)** Schematic of treatment timeline of MSK-BR-WGS-03.

**Fig S4**

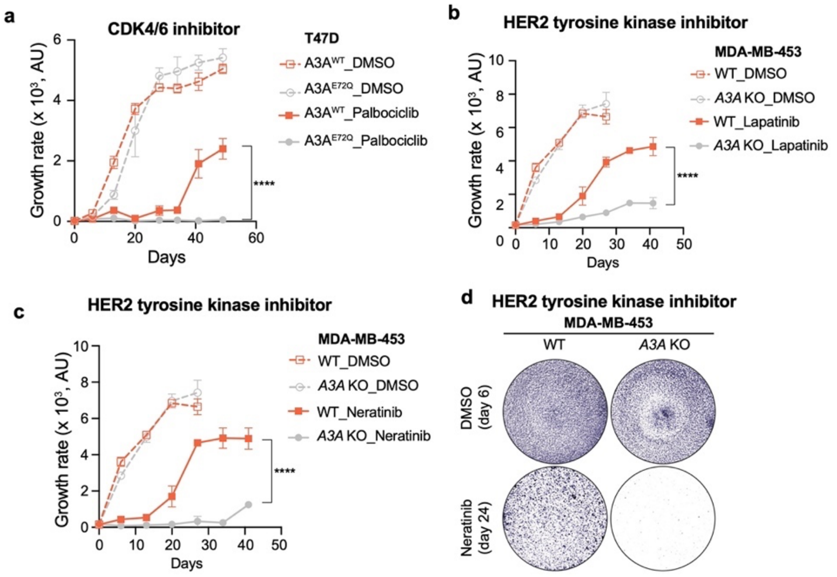


**Fig S4. APOBEC3 activity drives therapeutic resistance in breast cancers. (a)** Growth curves of T47D A3A$^{WT}$ and A3A$^{E72Q}$ cells treated with DMSO or palbociclib (500 nM). Data are represented as mean ± SD of three replicates. The groups were compared using two-way ANOVA test: **** $p$<0.0001. AU is abstract unit. **(b, c)** Growth curves of MDA-MB-453 WT and *A3A* KO cells treated with DMSO or **(b)** lapatinib (2 µM) or **(c)** neratinib (100 nM). Data are represented as mean ± SD of three replicates. The groups were compared using two-way ANOVA test: **** $p$<0.0001. **(d)** Crystal violet staining of MDA-MB-453 WT and *A3A* KO cells treated with DMSO (for 6 days) or neratinib (100 nM, for 24 days). Images are representative of $n$ = 3 replicates.

**Fig S5**

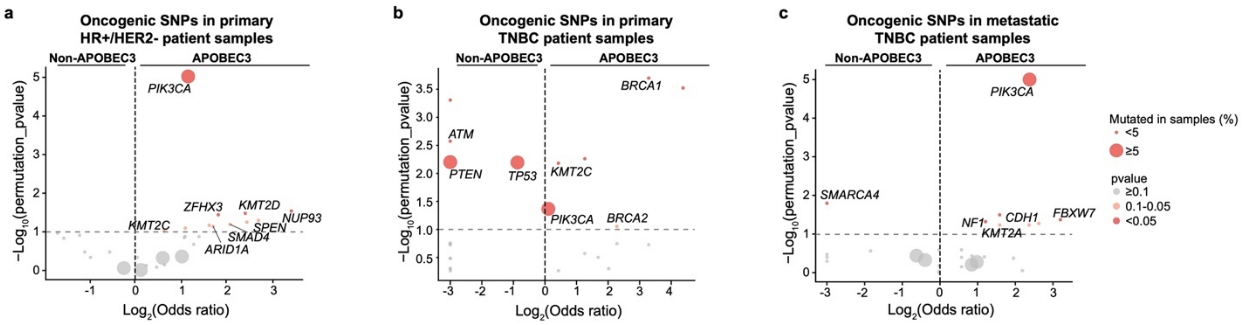


**Fig S5. Gene enrichment in breast cancer samples.** Volcano plots depicting enrichment of genes in primary HR+/HER2- and primary or metastatic TNBC samples categorized according to the dominant mutational signature.

### Fig S6

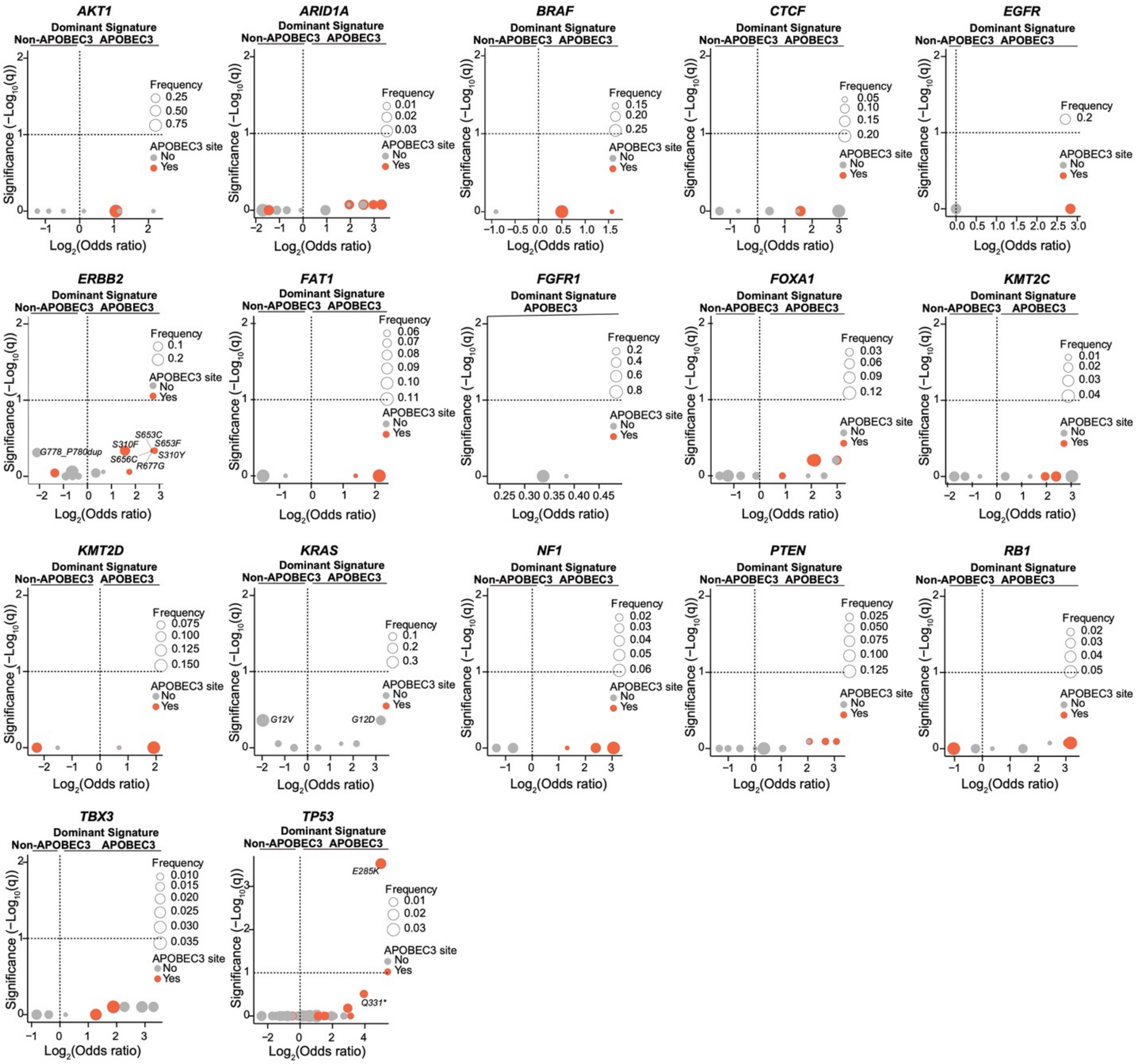


**Fig S6. Site-specific enrichment in HR+/HER2- breast cancer samples.** Volcano plots depicting site-specific enrichment of the indicated genes categorized according to the dominant mutational signature.

**Fig S7**

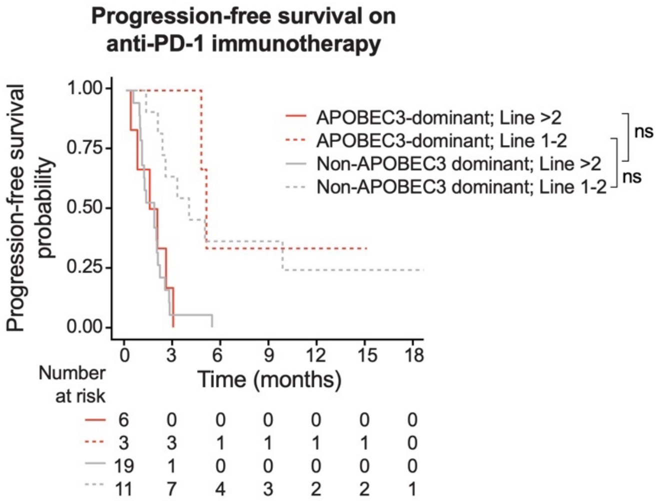


**Fig S7. Outcome of HR+/HER2- breast cancers to anti-PD-1 immunotherapy.** Kaplan-Meier curves displaying progression-free survival probability of patients with HR+/HER2- metastatic breast cancers treated with anti-PD-1 immunotherapy. The patients are categorized according to the dominant mutational signatures of the biopsy obtained prior to treatment start and the line of treatment. The groups were compared using the log-rank test: ns not significant.
